# Diet-induced MASH liver fibrosis promoted by EphB2 can be targeted by small molecule tetramerization inhibitors

**DOI:** 10.1101/2025.11.19.688968

**Authors:** Masaaki Yoshigi, Severin Donald Kamdem, Hanghang Wang, Amav Khambete, Francis Sprouse, Ernest M. Kameni, Estelle Atiogue, Sarah Pellizzari, Kimberley J. Evason, My N. Helms, Michael Goodman, Shuang Wang, Jisun So, Hyun Cheol Roh, Sihem Boudina, Mark Henkemeyer, Patrice N. Mimche

**Affiliations:** Department of Dermatology, Indiana University School of Medicine, Indianapolis IN 46202, USA; Department of Medicine, Division of Gastroenterology and Hepatology, Indiana University School of Medicine, Indianapolis, IN46202, USA; Department of Pathology, Division of Microbiology & Immunology, University of Utah, Salt Lake City, UT 84112, USA; Department of Neuroscience, Kent Waldrep Foundation Center for Basic Research on Nerve Growth and Regeneration, Peter O’Donnell Jr. Brain Institute, UT Southwestern Medical Center, Dallas, TX 75390, USA; Department of Pathology, Huntsman Cancer Institute, University of Utah, Salt Lake City, UT 84112, USA; Department of Internal Medicine, Division of Respiratory, Critical Care and Occupational Pulmonary Medicine, University of Utah, Salt Lake City, UT84132, USA; Department of Nutrition and Integrated Physiology, University of Utah, Salt Lake City, UT 84112, USA; Division of Liver Diseases, The Icahn School of Medicine at Mount Sinai, New York, NY 10029, USA; Department of Biochemistry and Molecular Biology, Indiana University School of Medicine, Indianapolis

## Abstract

The EphB2 receptor tyrosine kinase is thought to participate in numerous fibroinflammatory disorders. In metabolic dysfunction-associated steatohepatitis (MASH), we find EphB2 becomes strongly overexpressed and overactive in hepatic stellate cells (HSCs) from humans with the disease and from mice fed liver-injuring high fat diets. Genetic deletion of EphB2 or inactivation of its tyrosine kinase catalytic domain suppressed diet-induced MASH fibrosis, while a kinase overactive point mutant displayed exacerbated steatosis and hepatic damage. Silencing EphB2 in primary HSCs dampened the ability of TGF-β/SMAD signals to stimulate the transdifferentiation of stellate cells into profibrotic myofibroblasts, and HSC-specific deletion of the receptor, but not hepatocyte deletion, reduced liver scarring in multiple mouse models, even after fibrosis was established. Finally, a newly developed small molecule tetramerization inhibitor that targets EphB2-Ephrin receptor-ligand interactions effectively blunts inflammation and fibrosis in chemical and diet-induced liver injury models, demonstrating that therapeutically targeting EphB2 can counter MASH fibrosis.

## Introduction

Metabolic dysfunction-associated steatohepatitis (MASH) is a chronic inflammatory/ fibrotic disease that stems from progressive worsening of metabolic dysfunction-associated steatotic liver disease (MASLD). Viewed together, MASLD and MASH have become the most prevalent liver disease in developed nations due to the rise in type 2 diabetes and obesity^1,2^. In MASH, the progressive accumulation of liver fat (steatosis) and continual increasing liver inflammation leads to a chronic downward spiral of intensifying tissue fibrosis and hepatic injury that can advance to scarring, cirrhosis, and organ failure. Liver fibrosis in MASH is attributed to excessive activation of normally quiescent hepatic stellate cells (HSCs) by TGF-β/SMAD signals, which causes them to transdifferentiate into proliferating, profibrotic myofibroblasts that produce excessive collagen and other extracellular matrix (ECM) material characteristic of the disease^3,4^.

Hepatic fibrosis is associated with poor outcome in patients with MASH^5,6^. Despite ongoing progress in understanding the molecular mechanisms that drive MASH, treatments are limited to the recently FDA approved thyroid hormone receptor-β agonist and glucagon-like peptide-1 receptor agonists, both of which indirectly target fibrosis^7-9^. Additional mechanistic insight into MASH is needed to better target this disease.

Especially important will be the characterization of key membrane-anchored cell signaling pathways that work in the liver with TGF-β/SMAD to drive HSC activation and myofibroblast expansion to cause the downward spiral of increasing inflammation and fibrosis that leads to MASH. Single nucleus RNA-sequencing has recently identified potential cell-cell communication hubs critical for fibrosis in advanced MASH, including the receptor tyrosine kinase EphB2^10,11^, which we previously showed participates in liver fibrosis and scaring caused by infectious and chemical agents^12,13^.

EphB2 is one of fourteen highly related Eph molecules that comprise the largest family of transmembrane receptor tyrosine kinases (EphA1-A8, A10 and EphB1-B4, B6). The Eph receptors interact with Ephrin ligands, which are also tethered to the plasma membrane through either a glycosylphosphatidylinositol-linked moiety (EphrinA1-A5) or a transmembrane segment followed by a short intracellular tail (EphrinB1-B3)^14^. As both the Eph and Ephrin molecules are membrane localized, their interactions typically occur at sites of cell-cell contact and result in the activation of bidirectional signal transduction cascades into both the receptor-expressing cell (forward signaling) and ligand-expressing cell (reverse signaling)^15^. Prior literature suggests Eph-Ephrin interactions participate in diverse biological processes relevant to fibrogenesis^16^, including cell migration/proliferation, epithelial-to-mesenchymal-transition (EMT), angiogenesis, inflammation, wound healing, and tissue remodeling^17-20^. However, it remains poorly understood how EphB2 may participate in MASH fibrosis.

Here, we show that EphB2 expression and its activation are elevated in fibrotic livers from humans with MASH and in mice fed high fat fibrosis-inducing diets. We find EphB2 is required for the development of fibrosis and that this is driven by forward signaling as point mutation that inactivates the tyrosine kinase domain mitigated disease pathology, while point mutation that constitutively over-activates the kinase domain exacerbated liver scaring. Silencing EphB2 in hepatic stellate cells blocked their ability to become activated and transdifferentiate into ECM-producing myofibroblasts in response to TGF-β *in vitro*, and dampened liver fibrosis *in vivo*, as did treatment with novel small molecule Eph-Ephrin tetramerization inhibitors described in our accompanying manuscripts^21,22^. Overall, this work uncovers EphB2 forward signaling as a key mediator of HSCs activation by working with TGF-β/SMAD signaling to bring about liver fibrosis and scaring with potential implications for the treatment of MASH.

## Results

### EphB2 is up-regulated in human and mouse fibrotic MASH livers

To confirm if hepatic EphB2 expression correlates with the progression of fibrosis, we evaluated *EPHB2* mRNA levels in a publicly available transcriptomic dataset of patients with MASLD/MASH at various stages of fibrosis^23^. We found increased hepatic *EPHB2* transcripts at early stages of disease that intensified to reach very high levels of expression at fibrosis stage 4 (F4) in MASH (**Extended Data Fig. 1a**). Next, we stained liver biopsies of patients at different stage of fibrosis (F0-F4) with a phospho-EphB2 (pEphB2) specific antibody which is a readout of EphB2 signaling^22^, and found that receptor activation is strongly elevated in advanced MASH fibrosis F4 (**Extended Data Fig. 1b,c**). To ascertain that our findings in humans with MASH correlates with murine models of diet-induced MASH^24^, we analyzed an RNA sequencing dataset of livers from mice fed either a standard chow diet or the obesogenic Gubra Amylin NASH (GAN) diet^25^ for 20 weeks and found hepatic *Ephb2* transcripts to be very strongly up-regulated in GAN diet fed mice compared to chow fed mice (**Extended Data Fig. 1d**). We then assessed hepatic *Ephb2*, *Vim*, *Cd11b* mRNA levels in mice fed the GAN diet for 7, 12,18 and 26 weeks and found a significant progressive increase in mRNA expression of these genes, all reaching very high levels at week 26, with *Ephb2* transcripts increasing ∼75-fold compared to the levels detected in livers from normal chow fed mice (**Extended Data Fig. 1e**). Staining with the pEphB2 antibody showed much higher levels of activated receptor protein in liver sections of mice fed the GAN diet compared to chow fed mice (**Extended Data Fig. 1f**). Finally, we assessed hepatic EphB2 and αSMA protein levels in mice fed the non-obesogenic choline-deficient L-amino acid high fat diet (CDAA HFD) for 12 weeks and found the amount of both proteins increased compared to standard chow fed mice (**Extended Data Fig. 1g**). It was particularly noted that the level of EphB2 receptor protein was barely detectable in the liver lysates from normal chow fed mice, though it became strongly up-regulated in the livers from CDAA HFD fed mice. Taken together, this data shows hepatic EphB2 expression becomes strongly elevated in humans with MASH and in mice that have consumed a liver-injuring MASH diet.

### EphB2 deficiency mitigates diet-induced MASH fibrosis

To dissect the potential involvement of EphB2 in the development of MASH, we fed male *Ephb2^+/+^* wild-type (*WT*) and germline *Ephb2^-/-^* knockout (*KO*) littermate mice either a normal chow diet or the GAN diet for 26 weeks and assessed the level of hepatic steatosis, inflammation, and fibrosis. While the *Ephb2^-/-^ KO* mice are normal appearing and long-lived, we found they gained significantly less weight compared to the *Ephb2^+/+^ WT* controls when fed either diets during the 26 week period (**Extended Data Fig. 1h,i**), this despite an increased food intake in the mutant*s* (**Extended Data Fig. 1j**). Liver-to-body weight ratio was significantly reduced in GAN-fed *Ephb2^-/-^*compared to *Ephb2^+/+^* mice (**Extended Data Fig. 1k**), and glucose tolerance was significantly improved in the *KO* compared to *WT* (**Extended Data Fig. 1l**). These data indicate that EphB2 deficiency mitigates metabolic syndrome associated with diet-induced obesity in mice.

Histologic assessment of liver sections from *WT* mice fed the GAN diet for 26 weeks showed extensive steatosis (H&E staining) and collagen deposition (Masson’s Trichrome staining), whereas GAN diet fed *KO* mice had greatly reduced steatosis and collagen deposition (**Fig. 1a**). Plasma levels of alanine aminotransferase (ALT), a marker of liver damage, and total cholesterol were both significantly reduced in GAN diet fed *KO* mice compared to *WT*, with no statistical difference detected for plasma aspartate aminotransferase (AST) or triglycerides (**Fig. 1b,c)**. We performed bulk RNA-sequencing of the livers from GAN diet fed *KO* and *WT* animals and found that the top 25 genes significantly down-regulated in the *Ephb2-/-* mice were those implicated in fibrosis (e.g. *Col1a1, Col1a2, Col3a1, Vim, Col6a3,* and *Myof*) and inflammation (e.g *Clec7a, Cerk, Axl, Tlr4, H2-Q1, Lgals3,* and *Vcam1*) (**Fig. 1d**). In contrast, it was generally noted the livers from GAN fed *KO* retained high expression of genes involved in normal function. Additional analysis of the sequencing data revealed the signalling pathways significantly down-regulated in GAN diet fed *KO* mice were those involved in fibrosis and inflammation (**Extended Data Fig. 1m**). Quantitative PCR of selected fibrotic (*Col1a1, Timp1, Edn, Acta2, Pdgfrβ,* and *Vim*) and inflammatory (*Cd11b, Il-6, Tnf-α, Ccr2, Tgf-β1*, and *Tgf-β2*) markers confirmed all of these genes were significantly down-regulated in GAN diet fed *KO* mice (**Fig. 1e,f**). Immunophenotyping further showed a significant reduction in hepatic macrophage subsets (CD11b+F4/80+ and CD11b+Gr1+ cells) in the GAN fed *KO* mice, with no notable change in T (CD3) or B (CD19) cell populations (**Extended Data Fig. 2a,b**). Together, these data support the hypothesis that EphB2 directly participates in the pathogenesis of diet-induced MASH fibrosis with potential roles in bringing about steatosis, inflammation, and fibrosis.

**Fig. 1:**
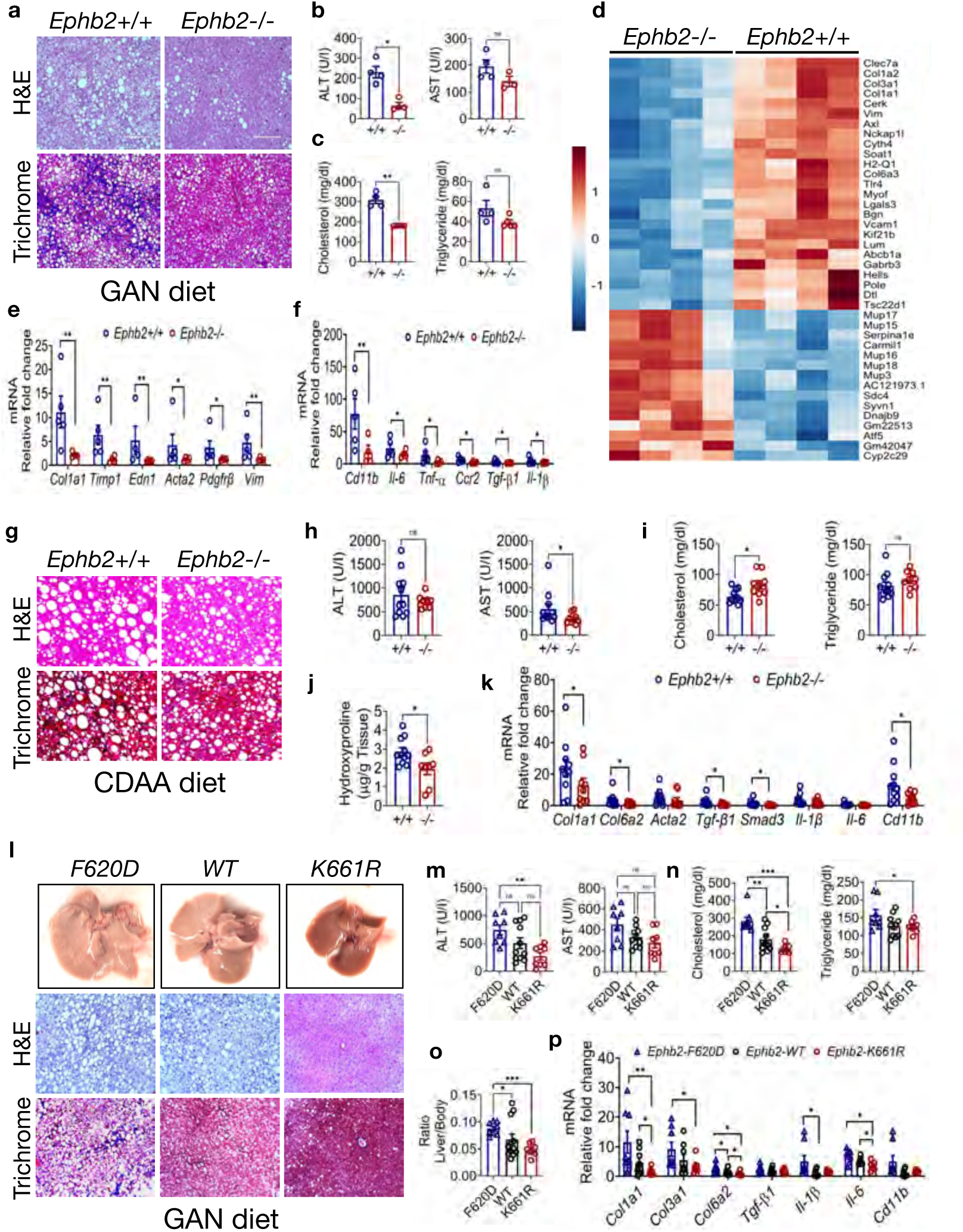
EphB2 tyrosine kinase signaling is needed for diet-induced MASH fibrosis. **a**, Representative images of liver sections from *Ephb2+/+* and *Ephb2-/-* mice fed the GAN diet for 26 weeks stained with *H&E* and Masson’s Trichrome. **b-c**, Plasma levels of liver damage enzymes ALT and AST (**b**) and cholesterol and triglyceride (**c**) from *Ephb2+/+* and *Ephb2-/-* mice fed the GAN diet for 26 weeks (*n* = 4 mice/group, \**p* < 0.05, \*\**p* < 0.01 by Student’s t test). **d**, Heatmap depicting the top 40 differentially expressed genes from bulk RNA sequencing of livers samples of *Ephb2+/+* and *Ephb2-/-* mice fed the GAN diet for 26 weeks. **e,f**, Liver mRNA levels of (**e**) fibrotic markers *Col1a1, Timp1*, *Edn1*, *Acta2*, *Pdgfrβ*, *Vim* and (**f**) inflammatory markers *Cd11b*, *Il-6*, *Tnf-α*, *Ccr2*, *Tgf-β1*, *Il-1β* from GAN diet fed *Ephb2+/+* and *Ephb2-/-* mice (*n* = 5 mice/group, \**p* < 0.05, \*\**p* < 0.01 by Student’s t test). **g**, Representative images of liver sections from *Ephb2+/+* and *Ephb2-/-* mice fed the CDAA HFD diet for 12 weeks stained with *H&E* and Masson’s Trichrome. **h-i**, Plasma levels of ALT and AST (**h**) and cholesterol and triglyceride (**i**) from CDAA HFD fed *Ephb2+/+* and *Ephb2-/-* mice (*n* = 9-11 mice/group, \**p* < 0.05, \*\**p* < 0.01 by Student’s t test. **j**, Level of hydroxyproline quantified in the livers from CDAA HFD fed *Ephb2+/+* and *Ephb2-/-* mice (*n* = 7-10 mice per group, \**p* < 0.05, \*\**p* < 0.01 by Student’s t test). **k**, Liver mRNA levels of fibrotic and inflammatory markers *Col1a1*, *Col6a2*, *Acta2*, *Tgf-β1*, *Smad3*, *Il-1β*, *Il-6*, *Cd11b* from CDAA HFD fed *Ephb2+/+* and *Ephb2-/-* mice (*n* = 9-11 mice/group, \**p* < 0.05, \*\**p* < 0.01 by Student’s t test). **l**, Representative images of livers from EphB2-F620D, wild type, and EphB2-K661R mice fed the GAN diet for 26 weeks photographed, and sections stained with *H&E* and Masson’s Trichrome. **m-o**, Plasma levels of (**m**) ALT and AST and (**n**) cholesterol and triglycerides, and (**o**) liver to body weight ratio from GAN fed EphB2-F620D, wild type, and EphB2-K661R mice (*n* = 8-11 mice per group, \**p* < 0.05, \*\**p* < 0.01, \*\*\**p* < 0.001 by ANOVA). **p**, Liver mRNA levels of fibrotic and inflammatory markers *Col1a1*, *Col3a1*, *Col6a2*, *Tgf-β1*, *Il-1β*, *Il-6*, *Cd11b* from GAN fed EphB2-F620D, wild type, and EphB2-K661R mice (*n* = 8-11 mice/group, \**p* < 0.05, \*\**p* < 0.01, by ANOVA). Data are presented as mean ± SEM, ns = not significant.

Given the systemic nature of the germline *Ephb2* deletion and the reduced body weight of the *KO* mice fed the GAN diet, it is possible that the above improvement of MASH pathology is related to the absence of obesity observed in these animals. We therefore assessed if EphB2 is required for the development of diet-induced hepatic fibrosis in a non-obese mouse model of MASH. Here, *WT* and *KO* littermates were fed the non-obesogenic choline deficient L-amino acid high fat diet (CDAA HFD)^26^ for 12 weeks. Histologic assessment of liver sections from these mice revealed no notable difference in the level of steatosis, although Masson’s Trichrome staining indicated collagen deposition was significantly diminished in the *KO* livers compared to *WT* (**Fig. 1g**). No significant differences in plasma levels of ALT or triglycerides were detected in CDAA fed, however plasma level AST was significantly reduced in the *KO* compared to *WT*, while plasma cholesterol was elevated in the mutants (**Fig. 1h,i**). Importantly, as expected, the *WT* mice fed the GAN diet showed ∼5-fold higher plasma cholesterol compared to the counterpart CDAA fed *WT* mice, while both diets resulted in elevated ALT/AST, indicative of liver injury, which was especially noted in the CDAA fed animals. Consistent with reduced collagen deposition detected in the Masson’s Trichrome stains of CDAA fed *KO* mice, hepatic hydroxyproline (a marker of collagen turnover) was also significantly reduced in the livers from the mutant mice compared to *WT* (**Fig. 1j**).

Quantitative PCR assessment of hepatic gene expression of selected fibrotic and inflammatory markers further showed a significant reduction of *Col1a1, Col6a2, Tgf-β1*, *Smad3*, and *Cd11b* transcripts in the livers of CDAA fed *KO* mice compared to *WT* (**Fig. 1k**), and immunophenotyping of selected hepatic myeloid cell subsets showed a significant reduction of inflammatory monocyte (Ly6C+) and Kupffer cells (F4/80+) in the mutants (**Extended Data Fig. 2c**). Altogether, the data show hepatic fibroinflammatory pathologies induced by either a GAN or CDAA HFD are strongly reduced when the mouse is globally deleted for EphB2.

### EphB2 forward signaling promotes MASH fibrosis

To determine if EphB2-mediated tyrosine kinase forward signaling is involved in diet-induced MASH fibrosis, mice with point mutations in key residues of the EphB2 catalytic domain *Ephb2^F620D/F620D^* (kinase-overactive, gain-of-function^27^), *Ephb2^K661R/K661R^* (kinase-dead, loss-of-function^28^), and *Ephb2^+/+^* (*WT*) littermate controls were fed the obesogenic GAN diet for 26 weeks. Compared to the *WT* animals, we found elevated steatosis and collagen deposition in liver sections from GAN diet fed *F620D* kinase-overactive mice, whereas the livers from *K661R* kinase-dead appeared like knockouts and showed much less steatosis and collagen deposition than the *WT* (**Fig. 1l**). While a significant change in the level of hepatic hydroxyproline was not observed in these mice (**Extended Data Fig. 1n**), the plasma levels of ALT and AST were both elevated in GAN diet fed *F620D* mice compared to the *WT* animals, and even more so when compared to the *K661R* mutants (significantly for ALT only) (**Fig. 1m**). Significant differences in plasma cholesterol and triglycerides was also detected, which were both elevated in the *F620D* mice compared to *WT* mice, and even more so when compared to the *K661R* mutants (**Fig. 1n**). Liver to body weight ratio was also significantly increased in the GAN fed *F620D* mice compared to *WT,* and even more so when compared to the *K661R* mutants (**Fig. 1o**). Assessment of mRNA levels of fibrotic (*Col1a1*, *Col3a1*, and *Col6a2*) and inflammatory (*Tgf-β1*, *Il-1β*, *Il-6*, and *Cd11b*) markers revealed that most were elevated in the livers from GAN fed *F620D* mice compared to *WT*, and even more so when compared to the *K661R* mutants (**Fig. 1p**). This data is consistent with the idea that EphB2-mediated tyrosine kinase forward signaling is a major driver of GAN diet-induced MASH.

Given the obesogenic nature of the GAN diet that can impact liver inflammation and fibrosis, we also examined the involvement of EphB2 tyrosine kinase forward signaling in the carbon tetrachloride (CCl_4_) model of hepatic fibrosis^12,13^. Here, *F620D* kinase-overactive, *K661R* kinase-dead, and *WT* littermate mice were injected with CCl_4_ (10%, 2 µl/g) three times a week for 4 weeks to induce liver injury, inflammation, and fibrosis. The resulting H&E, Masson’s Trichrome, and Sirius red stained histological sections all showed greatly increased damage and collagen accumulation in the livers of CCl_4_-injured *F620D* mice when compared to the *WT* counterparts, while the *K661R* mice phenocopied the *Ephb2-/-* knockout^12,13^ and showed hepatic tissue that appeared more like a normal uninjured liver (**Extended Data Fig. 3a**). Increased liver damage in the CCl_4_-injured *F620D* kinase-overactive mice was confirmed by detecting a significant increase in hepatic hydroxyproline when compared to the injured *WT* mice, while the hydroxyproline levels in CCl_4_-injured *K661R* kinase-dead mice were even lower and more significant than that of *WT* (**Extended Data Fig. 3b**). Plasma levels of ALT and AST were also elevated in CCl_4_-injured *F620D* mice when compared to the *WT* counterparts, and significantly higher when compared to the *K661R* kinase-dead mice (**Extended Data Fig. 3c**). An enhanced inflammatory response of EphB2 kinase-overactive mice following liver injury was documented by finding that hepatic CD11c+ (cDC1), CD11b+ macrophage, and F4/80 Kupffer cell populations were all significantly elevated in CCl_4_-injured *F620D* livers compared to the *WT*, and even more significantly elevated when compared to the K661R kinase-dead mice (**Extended Data Fig. 3d**).

Gene expression of fibrotic markers (*Col1a1*, *Col3a1*, *Acta2*, *Timp1*, *Col1a2* and *Col6a2*) further revealed that most were significantly elevated in the livers from the CCl_4_-injured *F620D* kinase-overactive mice when compared to *WT*, while the livers from *K661R* mice were protected as all markers showed significantly less expression when compared to the *F620D* mice, with *Col1a1*, *Col3a1*, *and Timp1* transcripts also significantly less than that detected in the injured *WT* livers (**Extended Data Fig. 3e**). Altogether, the above data indicates EphB2-mediated forward signaling facilitates hepatic inflammation and fibrosis following either diet-induced or chemical-induced injury to the liver.

### Expression of EphB2 in hepatocytes is not required for MASH fibrosis

To test if EphB2 expression in hepatocytes is needed for the development of MASH fibrosis, we generated conditional *Ephb2*^loxP/loxP^ mice and bred them with Albumin-*Cre* mice to selectively delete EphB2 expression in hepatocytes. Mice hemizygous for the hepatocyte-specific Cre driver and homozygous for *Ephb2*^loxP/loxP^ allele (*Hep^ΔEphb2^*) and Cre-negative *Ephb2*^loxP/loxP^ littermate controls (*Hep^WT^*) were fed the CDAA diet for 8 weeks. Histologic assessment of the resulting livers showed no difference in steatosis (H&E) or collagen deposition (Masson’s Trichrome) between the two groups (**Extended Data Fig. 4a**). Furthermore, other than a slightly elevated ALT, all other tested parameters of fibrosis or inflammation indicated the *Hep^ΔEphb2^* livers appeared no different from the *Hep^WT^* controls (**Extended Data Fig. 4b-h**). These data indicate deletion of EphB2 expression in hepatocytes does not alter the progression of MASH fibrosis in CDAA fed mice.

### TGF-β induced EPHB2 expression in human HSCs is SMAD-dependent

To identify the main hepatic cell type that upregulates EphB2 expression in MASH fibrosis, we analyzed a publicly available single-nuclei RNA sequencing dataset of mice with advanced disease^10^. The data showed *Ephb2* transcripts were mostly elevated in HSCs, while those for one of its transmembrane ligands, EphrinB2 (*Efnb2*), were mainly found in endothelial cells which make extensive contact with stellate cells and communicate important cell-cell signals involved in hepatic inflammation and fibrosis (**Extended Data Fig. 5a**). We next used immunofluorescence to assess for activated, tyrosine phosphorylated EphB2 (pEphB2) receptor protein and αSMA (a marker of activated HSCs/myofibroblasts) in sections of human liver biopsies from healthy and advanced MASH fibrosis patients. We found that compared to the healthy liver, pEphB2 was strongly up-regulated in human advanced MASH and colocalized with high levels of αSMA (**Fig. 2a**). This indicates HSCs and their activated profibrotic myofibroblasts as the main hepatic cell types that upregulate EphB2 protein to potentiate its signaling in MASH.

**Fig. 2:**
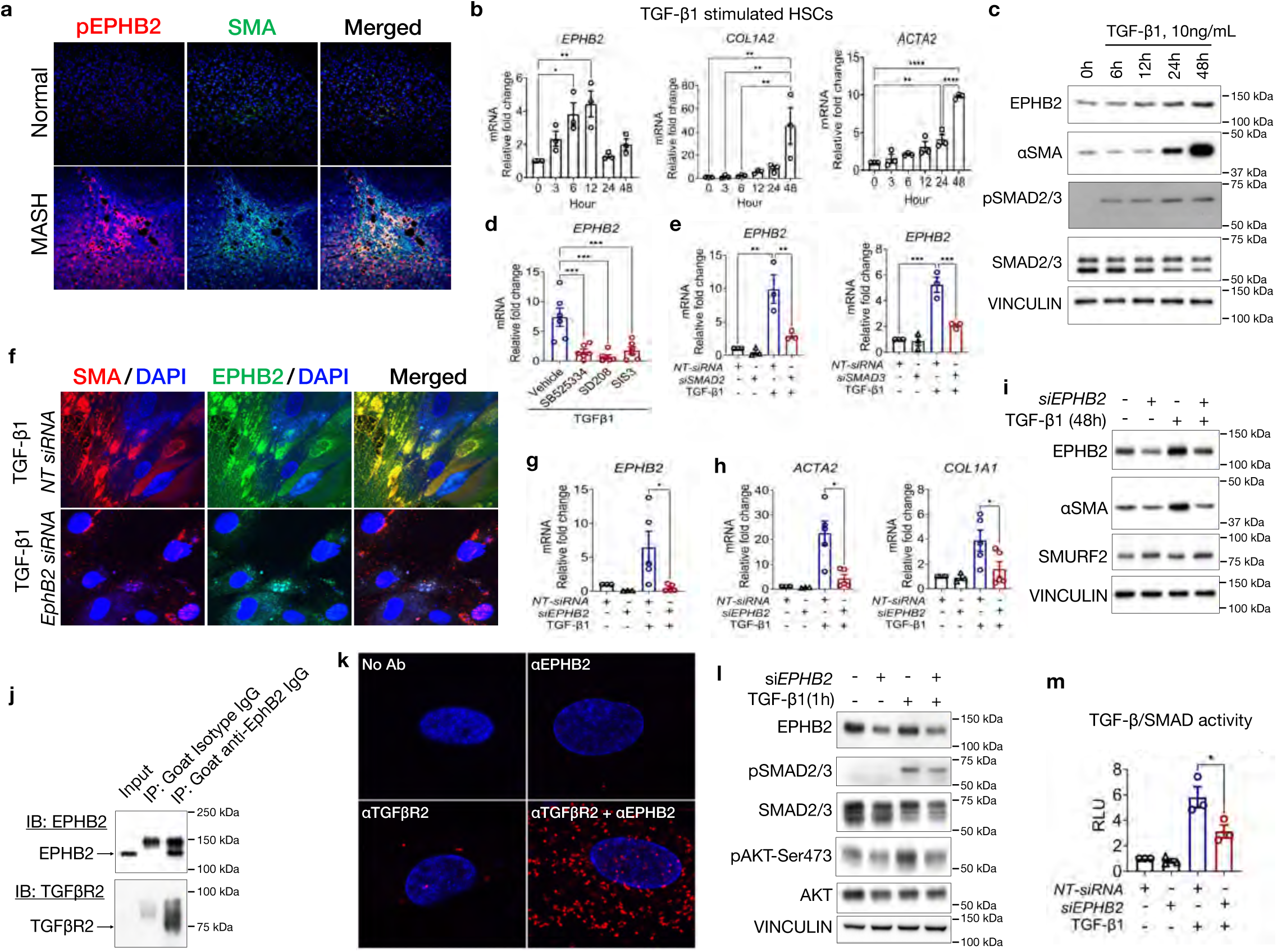
TGF-β/SMAD signaling in human hepatic stellate cells induces EPHB2 expression required for their trans-differentiation into activated myofibroblasts. **a**, Representative immunofluorescence images of human liver sections of healthy and MASH patients stained with antibodies that detect the active, tyrosine phosphorylated pEPHB2 protein (red) and αSMA (green), with DAPI counterstain for nuclei (blue). **b**, The mRNA levels of *EPHB2*, *COL1A2*, and *ACTA2* in TGF-β1 stimulated primary human HSCs at various time points (n=3). **c**, Immunoblot analysis for protein levels of EPHB2, aSMA, pSMAD2/3, SMAD2/3, and VINCULIN in TGF-β1 stimulated primary human HSCs at various time points. **d**, The mRNA levels of *EPHB2* upon inhibition of canonical TGF-β/SMAD signaling with small molecule TGFβR1 inhibitors (SB525334 and SD208) and SMAD3 inhibitor (SIS3) in TGF-β1 stimulated primary human HSCs (n=5-7). **e**, Effects of silencing SMAD2 and SMAD3 with Mission®-esiRNA on TGF-β-induced *EPHB2* mRNA expression in primary human HSCs (n=3). Non-targeting Mission®-esiRNA was used as control. **f**, Representative immunofluorescence images of TGF-β1 stimulated primary human HSCs transfected with *EPHB2*-siRNA and non-targeted control NT-siRNA, stained with antibodies against EPHB2 (green) and αSMA (red), with DAPI counterstain for nuclei (blue). **g**, The mRNA levels of *EPHB2* and (**h**) *ACTA2* and *COL1A1* from different HSCs lines treated with Mission® *EPHB2*-esiRNA (SIGMA), stimulated with TGF-β1 at 10 ng/ml, and normalized to the same HSCs lines treated with non-targeting Mission®-esiRNA (n=5). **i**, Immunoblot analysis for protein levels of EPHB2, αSMA, SMURF2, and VINCULIN in primary human HSCs stimulated with 10 ng/ml TGF-β1 for 48 hr showing the effects of silencing *EPHB2* with Mission®-esiRNA. **j**, Protein-protein interaction between EPHB2 and TGFβR2 detected by immunoprecipitation of EPHB2 in TGF-β1 stimulated human HSCs followed by immunoblot for EPHB2 and TGFβR2. **k**, Proximity ligation assay shows EPHB2 and TGFβR2 proteins are closely localized in TGF-β1 stimulated HSCs. **l**, Immunoblot analysis shows the effects of silencing *EPHB2* with Mission®-esiRNA on the expression of EPHB2, pSMAD2/3, SMAD2/3, pAKT-Ser473, AKT, and VINCULIN proteins in primary human HSCs stimulated with TGF-β1 for 1 hour. **m**, EPHB2-silenced and control primary human HSCs also transduced with an AAV8 expressing luciferase vector under the control of the TGFβ-SBE-promoter reporter were stimulated for 3 hr with TGF- β1 and then SMAD transcriptional activation was assessed by measuring luciferase activity (n=3). Data is presented as mean ± SEM, \**p* < 0.05, \*\**p* < 0.01, \*\*\**p* < 0.001, \*\*\*\**p* < 0.0001 by ANOVA, ns = not significant.

TGF-β cytokines are thought to be the main driver of MASH by activating the quiescent HSCs to transdifferentiate into ECM-producing myofibroblasts^29-33^. We therefore sought to determine if TGF-β signaling impacts EphB2 expression in cultured primary human HSCs. We found stimulation of HSCs in a time course up to 48 hr with recombinant human TGF-β1 significantly up-regulated *EPHB2*, *ACTA2*, and *COL1A2* mRNA levels (**Fig. 2b**) as well as EPHB2, αSMA and pSMAD2/3 protein levels (**Fig. 2c**). Not limited to TGF-β1, recombinant human TGF-β2 and TGF-β3 also up-regulated *EPHB2* and *ACTA2* mRNA and protein in primary human HSCs (**Extended Data Fig. 5b-e**). We next used selective pharmacological inhibitors^34^ of TGF-β receptor type I (TGFβR1, SD208 and SB525334) and SMAD3 (SIS3), finding that the increased expression of EPHB2 in TGF-β1 stimulated HSCs was dependent on TGF-β/SMAD signaling (**Fig. 2d**). Moreover, knockdown of *SMAD2* and *SMAD3* using siRNA also blocked TGF-β1-mediated upregulation of *EPHB2* mRNA in HSCs (**Fig. 2e**). We next analyzed a publicly available SMAD3 ChIP-sequencing dataset of TGF-β1 stimulated HSCs cell line^35^ and found SMAD binding elements (SBE) are present in the human *EPHB2* promoter region (**Extended Data Fig. 5f**). To validate the presence of SBE in the peak/promoter region of *EPHB2*, we performed a targeted SMAD3 ChIP-qPCR in control and TGF-β1 stimulated primary human HSCs and found that indeed SMAD3 occupied the *EPHB2* promoter region analyzed (**Extended Data Fig. 5g**).

### EPHB2 is required for HSCs activation

Given the above data that shows genetic deletion of EphB2 or point mutant inactivation of its tyrosine kinase catalytic domain mitigated HFD-induced MASH fibrosis and that EPHB2 becomes strongly expressed in TGF-β activated human HSCs, we reasoned that this receptor could be critical for hepatic stellate cell-to-myofibroblast transdifferentiation^13^. To test this idea, we silenced *EPHB2* in primary human HSCs with siRNA and then stimulated with TGF-β1 for 48 hr. The data showed that compared to the HSCs that received the control non-targeting siRNA, silencing *EPHB2* in HSCs strongly reduced expression of both the receptor protein (**Fig. 2f**) and mRNA (**Fig. 2g**). Moreover, silencing *EPHB2* suppressed the ability of TGF-β1 to stimulate HSC transdifferentiation into myofibroblasts as expression of *ACTA2* and *COL1A1* mRNA and αSMA protein levels were significantly reduced compared to the control non-targeting siRNA cells, and more resembled the levels detected in cells not exposed to the cytokine (**Fig. 2h,i**). We also noticed that silencing *EPHB2* in HSCs increased SMURF2 protein (**Fig. 2i**) as well as the mRNA levels of inhibitory SMADs (*SMAD7*, *SMURF1* and *SMURF2*) (**Extended Data Fig. 6a**) known to dampen TGF-β/SMAD signaling^36^.

Because EPHB2 is a transmembrane receptor tyrosine kinase, we reasoned the profibrotic activity of this protein could involve a potential interaction with TGF-β receptors. To address this possibility, we immunoprecipitated EPHB2 from primary human HSCs and found that TGFβR2 co-precipitated (**Fig. 2j**). Proximity ligation assay confirmed that EPHB2 is indeed in the vicinity of TGFβR2 (**Fig. 2k**), lending support for a potential crosstalk between EPHB2 and TGFβR2 in the regulation of TGF-β/SMAD signaling in fibrogenesis. To further confirm a role of EPHB2 in TGF-β/SMAD signaling, silencing *EPHB2* in HSCs stimulated with TGF-β1 for 1 hr attenuated pSMAD2/3 and pAKT (**Fig. 2l**), which are required for myofibroblast activation^37^. Furthermore, we transduced primary human HSCs with a lentivirus expressing luciferase under the control of a multimerized SBE and determined that TGF-β1-stimulated promoter activity was significantly reduced when *EPHB2* was silenced compared to the controls (**Fig. 2m**). The data above solidifies an interconnection between EPHB2 and TGFβR2, with the two receptors working together to potentiate each other’s expression/activity and to bring about the activation of HSCs.

Bulk RNA sequencing was then performed to gain insight in the signaling pathways that are affected when *EPHB2* is silenced in HSCs stimulated with TGF-β1. This revealed the top down-regulated genes in *EPHB2* silenced HSCs were involved in fibrosis including *EPHB2*, *DES*, *ITGA7*, *TIMP3*, *THBS2*, *ISLR*, *CILP*, *ASS1*, *RASSF2*, and *PDLIM3* (**Extended Data Fig. 6b,c**) which were validated by qPCR (**Extended Data Fig. 6d**). To identify the kinases that are altered when *EPHB2* is silenced, we performed a global kinome mapping and found that CDKs, CDKLs, and MAPKs serine/ threonine kinases were the main kinases dysregulated compared to TGF-β-stimulated control HSCs (**Extended Data Fig. 7**). Finally, as stellate cells actively migrate to injured areas of the liver to further spread ECM deposition and fibrosis, we subjected *EPHB2* silenced HSCs to wound healing assays and found significantly dampened migration compared to the control cells (**Extended Data Fig. 8**). Altogether, the above data point to a critical interplay of EPHB2 and TGF-β/SMAD signaling that can affect biological processes important in MASH, such as stellate cell transdifferentiation and migration.

### EphB2 deficiency in HSCs dampens MASH fibrosis

To better understand how EphB2 drives MASH fibrosis, *Ephb2*^loxP/loxP^ mice were bred with tamoxifen-inducible *PdgfrβCre*^ERT2^ mice to delete its expression in HSCs^38^. Mice hemizygous for the *PdgfrβCre*^ERT2^ Cre driver and homozygous for *Ephb2*^loxP/loxP^ (*HSC^ΔEphb2^*) and Cre-negative *Ephb2*^loxP/loxP^ littermates (*HSC^WT^*) were administered tamoxifen by intraperitoneal (i.p.) injection for 5 days and then fed the CDAA diet for 8 weeks. Histologic assessment of resulting liver sections showed no obvious difference in the level of steatosis (H&E), however, collagen deposition (Masson’s Trichrome) was clearly diminished in the CDAA fed *HSC^ΔEphb2^* animals compared to the *HSC^WT^* controls (**Fig. 3a**). While no differences in the plasma levels of ALT, AST, cholesterol, or triglycerides were observed in CDAA fed *HSC^ΔEphb2^* versus *HSC^WT^* mice (**Fig. 3b,c**), the reduced collagen deposition was confirmed by finding a significant decrease in hepatic hydroxyproline in the CDAA fed *HSC^ΔEphb2^* mice (**Fig. 3d**). Quantitative PCR of hepatic gene expression showed a significant reduction of *Col1a1 and Col3a1* mRNA levels in the livers of in the CDAA fed *HSC^ΔEphb2^* mice compared to the *HSC^WT^* controls (**Fig. 3e**), though no difference in inflammatory markers or myeloid cell populations were detected (**Extended Data Fig. 9a,b**). Bulk RNA sequencing further revealed that some of the top up-regulated genes in the CDAA fed *HSC^WT^* livers that were down-regulated in *HSC^ΔEphb2^* livers included *Lrg1*, *Egfr*, *Fgf1*, *Orm*1, *Lbp*, *Serpina3n*, *Saa2*, *Saa1*, and *Nt5e,* which are all known to be up-regulated in human MASH (**Extended Data Fig. 9c**)^10^. Pathway analysis indicated that biological processes implicated in fibrogenesis were down-regulated in CDAA fed *HSC^ΔEphb2^* compared to *HSC^WT^* mice (**Extended Data Fig. 9d**).

**Fig. 3:**
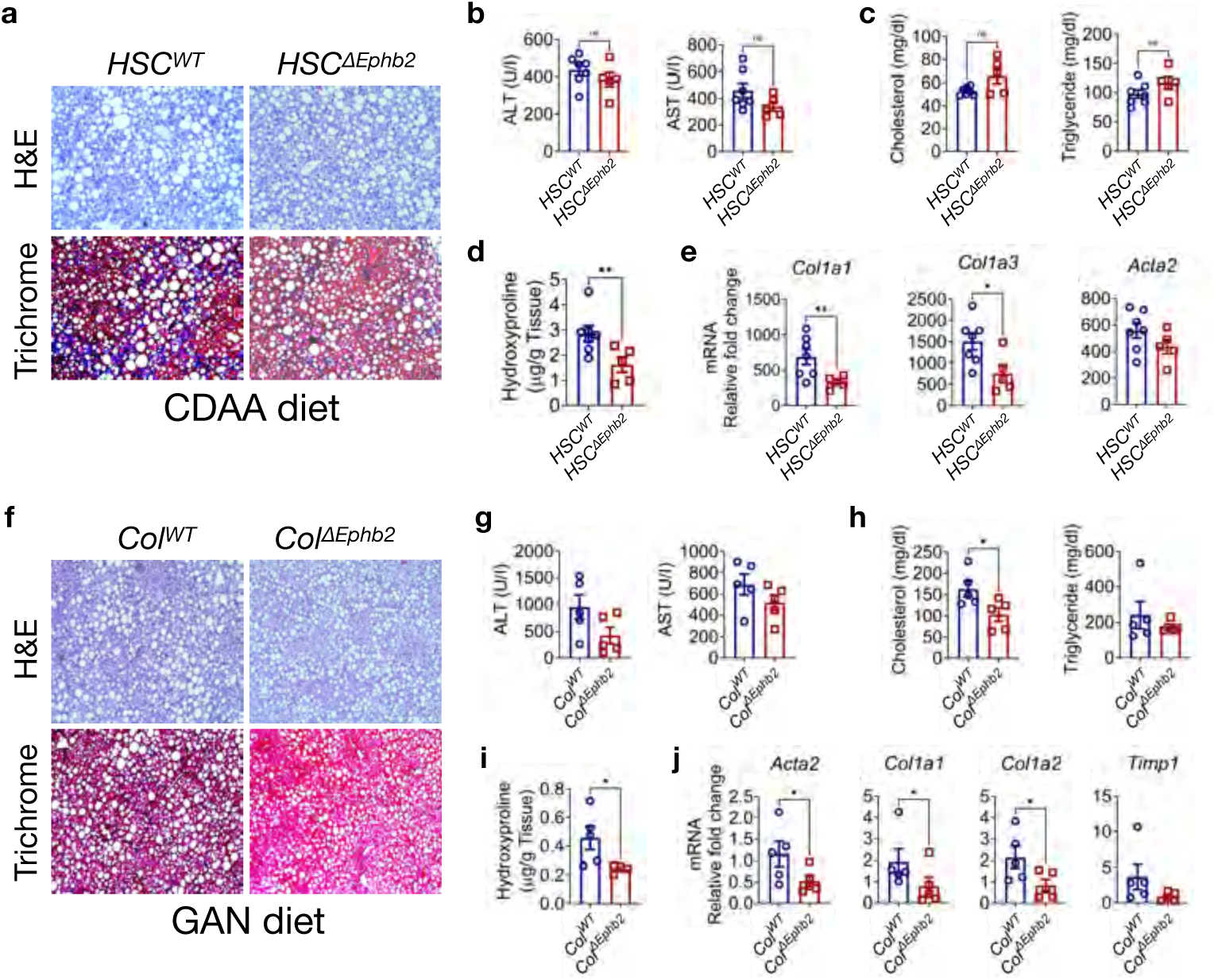
Deletion of *Ephb2* in liver mesenchymal cells mitigates CDAA and GAN diet-induced MASH fibrosis. **a-e**, Mice hemizygous for a *PdgfrβCre^ERT2^* Cre driver and homozygous for a loxP-flanked conditional *Ephb2^loxP/loxP^* allele were repeated injected with tamoxifen to selectively delete *EphB2* expression in hepatic stellate cells, and then the animals were fed the CDAA HFD diet for 8 weeks. **a**, Representative images of resulting liver sections from Cre-negative, tamoxifen-injected *Ephb2^loxP/loxP^* control mice (*HSC^WT^*) and Cre-containing, tamoxifen-injected *Ephb2^loxP/loxP^* experimental mice (*HSC^ΔEphb2^*) stained with *H&E* and Masson’s Trichrome. **b-c**, Plasma levels of (**b**) liver damage enzymes ALT and AST and (**c**) cholesterol and triglycerides from CDAA fed *HSC^WT^* and *HSC^ΔEphb2^* mice. **d**, Hydroxyproline quantified from livers of CDAA fed *HSC^WT^* and *HSC^ΔEphb2^* mice. **e**, Liver mRNA levels of *Col1a1*, *Col3a1*, *Col6a2*, *Acta2* from CDAA fed *HSC^WT^* and *HSC^ΔEphb2^* mice. Data are presented as mean ± SEM, \**p* < 0.05, \*\**p* < 0.01 by Student t test (*n* = 5-7 mice/group). **f-j**, Mice hemizygous for a *Col1a2Cre^ERT2^* Cre driver and either *Ephb2^+/+^* WT or homozygous for a loxP-flanked conditional *Ephb2^loxP/loxP^* allele were repeated injected with tamoxifen to selectively delete *EphB2* expression in hepatic stellate cells, and then the animals were fed the GAN HFD diet for 26 weeks. **f**, Representative images of resulting liver sections from Cre-containing, tamoxifen-injected *Ephb2^+/+^* control mice (*Col^WT^*) and Cre-containing, tamoxifen-injected *Ephb2^loxP/loxP^* experimental mice (*Col^ΔEphb2^*) stained with *H&E* and Masson’s Trichrome. **g-h**, Plasma levels of (**g**) ALT and AST, and (**h**) cholesterol and triglycerides in *Col^WT^* and *Col^ΔEphb2^* mice fed the GAN diet. **i**, Hydroxyproline quantified from the livers of *Col^WT^* and *Col^ΔEphb2^* mice fed the GAN diet. **j**, Liver mRNA levels of *Acta2*, *Col1a1*, *Col1a2*, *Timp1* in *Col^WT^* and *Col^ΔEphb2^* mice fed the GAN diet. Data is presented as mean ± SEM, \**p* < 0.05 by Student t test, ns = not significant (*n* = 5 mice/ group).

Given that *Col1a2* was one of the top up-regulated genes in *WT* mice fed the GAN diet (**Fig. 1d**), we investigated the impact of specifically deleting EphB2 in Col1a2 expressing HSCs. We bred conditional *Ephb2*^loxP/loxP^ animals with mice that carry the tamoxifen-inducible *Col1a2Cre*^ERT2^ allele to delete EphB2 expression in HSCs. Mice hemizygous for the *Col1a2Cre*^ERT2^ Cre driver and homozygous for *Ephb2*^loxP/loxP^ (*Col^ΔEphb2^*) and control Cre-negative *Ephb2*^loxP/loxP^ (*Col^WT^*) littermates were administered tamoxifen i.p. for 5 days and then fed the GAN diet for 22 weeks. Histologic assessment of liver sections showed no obvious difference in the level steatosis (H&E) between the two groups, however, collagen deposition (Masson’s Trichrome) was clearly diminished in the livers of GAN diet fed *Col^ΔEphb2^* mice compared to the *Col^WT^* controls (**Fig. 3f**). While we did not observe a significant difference in the plasma levels of ALT, AST or triglycerides in GAN fed *Col^ΔEphb2^* compared to *Col^WT^* mice, plasma level cholesterol was significantly reduced in GAN fed *Col^ΔEphb2^* compared to *Col^WT^* mice (**Fig. 3g,h**).

Consistent with the Masson’s Trichrome histology, hepatic hydroxyproline content was significantly reduced in livers of GAN fed *Col^ΔEphb2^* mice compared to *Col^WT^* mice (**Fig. 3i**), though there was no significant difference in the liver to body weight ratio between them (**Extended Data Fig. 10a**). Fibrotic marker mRNA levels *Acta2, Col1a1, and Col1a2* were all significantly reduced in livers from GAN fed *Col^ΔEphb2^* mice compared to the *Col^WT^* counterparts (**Fig. 3j**), while no difference was detected for mRNA levels of inflammatory markers (**Extended Data Fig. 10b**). Most hepatic myeloid cell subsets did not show significant differences, except the cDC1 and Ly6C+ cell populations were significantly reduced in the *Col^ΔEphb2^* mice (**Extended Data Fig. 10c**). Taken together, these data emphasize the role of EphB2 in HSC/myofibroblast activation *in vivo* as conditional deletion of EphB2 using either *PdgfrβCre*^ERT2^ or C*ol1a2Cre*^ERT2^ drivers did not mitigate steatosis, though both approaches clearly suppressed HFD-induced fibrosis.

### EphB2 deficiency in HSCs attenuates CCl_4_-induced hepatic fibrosis

To determine if EphB2 expression in HSCs also contributes to chemically-induced hepatic fibrosis, a group of *Col^ΔEphb2^* and *Col^WT^* littermates were administered tamoxifen i.p. for 5 consecutive days and then treated with CCl_4_ three times a week for 4 weeks. Histologic assessment showed a reduction in collagen accumulation in sections of CCl_4_-injured livers from *Col^ΔEphb2^* mice compared to the *Col^WT^* mice (**Extended Data Fig. 11a**), that was associated with a significant reduction in hepatic hydroxyproline content (**Extended Data Fig. 11b**). While, no significant difference in the plasma ALT or AST was observed between the two groups (**Extended Data Fig. 11c**), hepatic mRNA levels of *Col1a1*, *Fn1*, *Acta2*, and *Timp1* were all significantly attenuated in the CCl_4_-injured *Col^ΔEphb2^* livers compared to the *Col^WT^* counterparts (**Extended Data Fig. 11d**). However, no difference was detected for expression of hepatic inflammatory markers (**Extended Data Fig. 11e**), consistent with immunophenotyping that indicated no significant difference in the levels of myeloid cell populations (**Extended Data Fig. 11f**). These data solidify the notion that EphB2 expression in HSCs participates in the profibrotic transdifferentiation of stellate cells into myofibroblasts, a major prerequisite for the development of hepatic fibrosis.

### Deletion of EphB2 in HSCs promotes regression of establish fibrosis

We next investigated whether deletion of EphB2 would promote the regression of established fibrosis. Here, male *Col^WT^* and *Col^ΔEphb2^* mice (6 to 8 weeks of age) were fed the non-obesogenic CDAA diet for a total of 12 weeks to induce steatosis, inflammation, and fibrosis. At week 8, after MASH fibrosis was established, animals were i.p. injected with tamoxifen each day for 1 week to delete EphB2 expression from the HSC/ myofibroblast compartment, and then the mice were allowed to continue the CDAA diet for an additional 3 weeks (**Fig. 4a**). Histologic assessment of resulting liver sections showed a strong reduction of collagen accumulation in the animals deleted for EphB2 (**Fig. 4b**), that was associated with a significant reduction in hepatic hydroxyproline content (**Fig. 4c**). However, no obvious difference in ALT, AST, cholesterol, or triglycerides was observed between the CDAA fed *Col^WT^* control and *Col^ΔEphb2^* mice (**Fig. 4d,e)**, even though the total hepatic mRNA level of *Ephb2* was significantly reduced in the HSC EphB2-deleted animals (**Fig. 4f**). Consistent with reduced collagen and hydroxyproline, hepatic gene expression of fibrotic markers *Acta2*, *Col1a1*, *Col1a2*, *Col3a1, Col6a2*, and *Pdgfrβ* were all significantly attenuated in the CDAA fed *Col^ΔEphb2^* EphB2-deleted mice compared to the *Col^WT^* controls (**Fig. 4g**). In contrast, no significant difference was detected for liver inflammatory markers (**Extended Data Fig. 12**). Taken together, these data show deletion of EphB2 can promote the regression of fibrosis when MASH is already established.

**Fig. 4:**
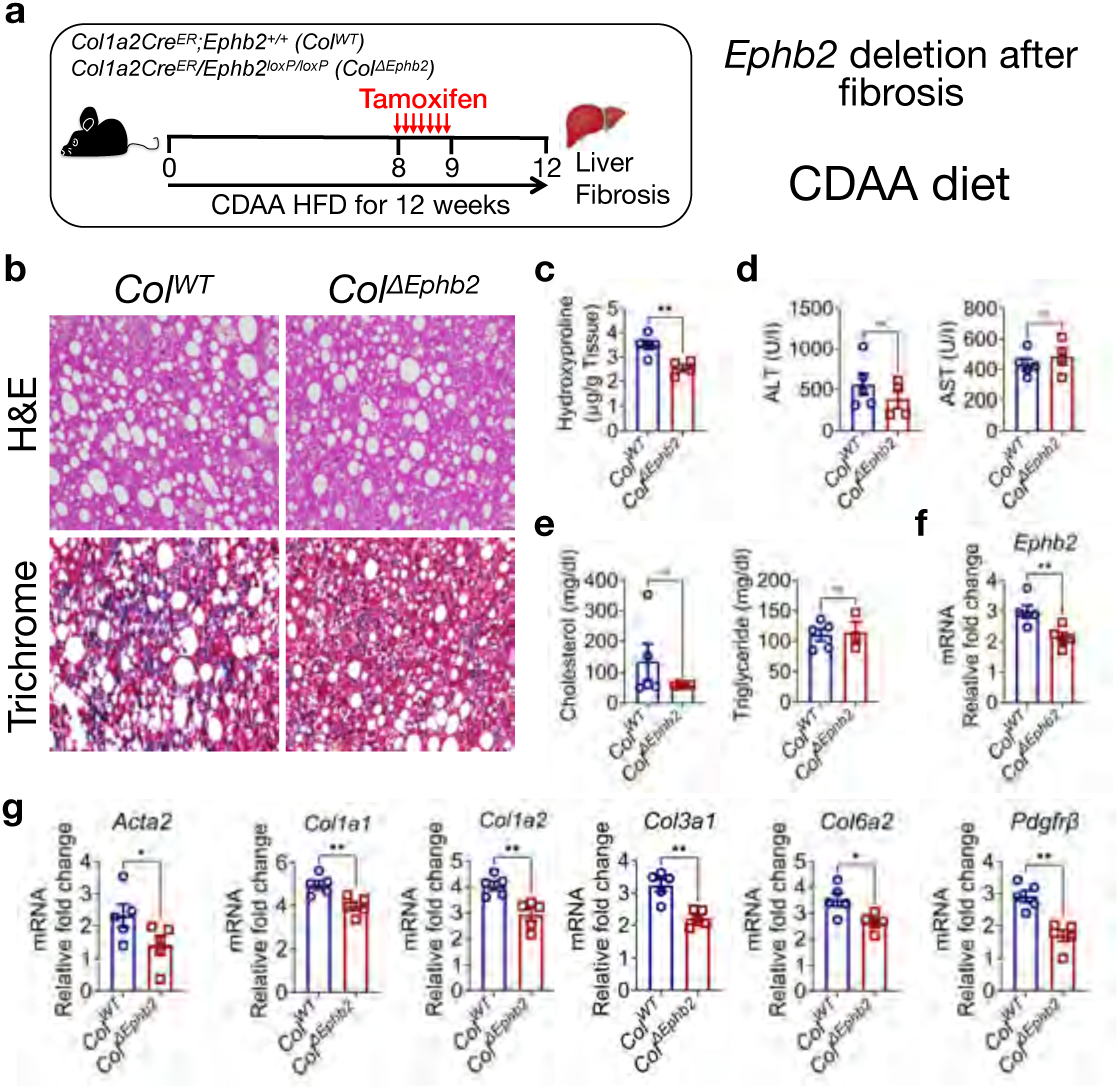
Deletion of *Ephb2* in liver mesenchymal cells when fibrosis is already established attenuates CDAA diet-induced MASH fibrosis. **a**, Experimental design used to delete EphB2 in liver *Col1a2*-expressing cells at week 8 when fibrosis is already present in *Col^WT^* and *Col^ΔEphb2^* mice fed the CDAA HFD for a total of 12 weeks. **b**, Representative images of liver sections of *Col^WT^* and *Col^ΔEphb2^* mice stained with *H&E* and Masson’s Trichrome to assess tissue damage and collagen deposition. **c**, Hydroxyproline quantified in the livers of *Col^WT^* and *Col^ΔEphb2^* mice (n = 4-5 mice/group). **d-e**, Plasma levels of (**d**) ALT and AST and (**e)** cholesterol and triglycerides from *Col^WT^* and *Col^ΔEphb2^* mice (*n* = 4-5 mice/group). **f**, Liver mRNA level of Ephb2 in *Col^WT^* and *Col^ΔEphb2^* mice (*n* = 5 mice/group). **g**, Liver mRNA levels of *Acta2*, *Col1a1*, *Col1a2*, *Col3a1*, *Col6a2*, *Pdgfrβ* in *Col^WT^* and *Col^ΔEphb2^* mice (*n* = 5 mice/group). Data is presented as mean ± SEM, \**p* < 0.05, \*\**p* < 0.01 by Student t test, ns = not significant.

### Eph-Ephrin tetramerization inhibitor A20 mitigates hepatic fibrosis in CCl_4_-injured mice

In the accompanying manuscripts^21,22^, we describe our discovery of new, first-in-class small molecule Eph-Ephrin tetramerization inhibitors that antagonize with submicromolar activity the ability of EphB2 to bind its EphrinB ligands with and form the high-affinity circular tetramer macromolecular structure necessary for these molecules to become activated to begin transducing their cell-cell bidirectional phosphotyrosine signals. To determine if our lead compound termed A20 could counter fibrotic signaling *in vitro*, primary human HSCs were stimulated for 48 hr with either TGF-β1, TGF-β2, or TGF-β3 to activate their transdifferentiation into profibrotic myofibroblasts, and then cultured for an additional 24 hr with the indicated TGF-β in the presence or absence of 5 µM of A20. This resulted in a strong and significant reduction in *COL1A1, COL1A2,* and *ACTA2* mRNA levels in A20-treated stellate cells compared to the no compound controls, no matter which TGF-β isoform was used (**Fig. 5a, Extended Data Fig. 13a,b**). A20 did not affect HSC cell viability, which is consistent with MMT survival studies presented in our accompanying manuscript^39^ that show the compound at up to 5 µM is well tolerated in pancreatic stellate cells and pancreatic cancer cells during 72 hr incubation.

**Fig. 5:**
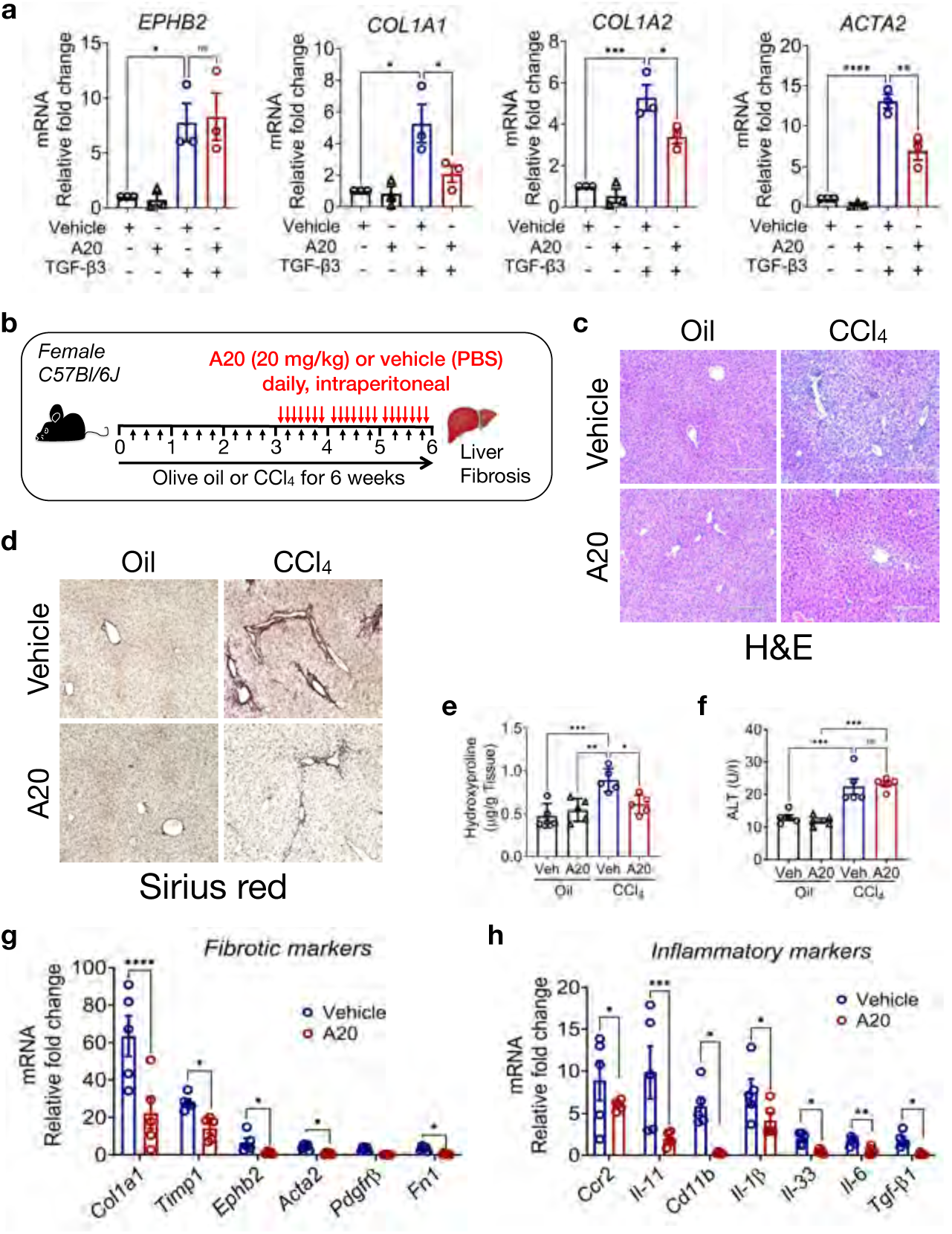
Tetramerization inhibitor A20 blunts TGFβ activation of human liver stellate cells in *vitro* and reduces CCl_4_-induced liver fibrosis in mice *in vivo*. **a**, Human primary HSC cells were first stimulated with 10 ng/ml TGF-β3 for 48 hr and then further incubated in the presence or absence of 5 µM of A20 tetramerization inhibitor compound (2x HCl salt) for an additional 24h. RNA was extracted and subjected to quantitative PCR to assess gene expression of profibrotic markers *EPHB2*, *COL1A1*, *COL1A2*, and *ACTA2*. Data represent Mean ± SEM (n=3/group \**p* < 0.05, \*\**p*< 0.01, \*\*\**p*<0.001, \*\*\*\**p*<0.0001 by ANOVA). **b**, Study outline for CCl_4_ fibrosis model in which female C57BL6/J mice were administered via intraperitoneal (i.p.) injection 10% CCl_4_ diluted in olive oil thrice weekly at a dose 2 µl/g for 6 weeks. Three weeks after starting CCl_4_ injections, either vehicle or A20 tetramerization inhibitor (a 2xHCl salt of A20 was dissolved in PBS and dosed at 20 mg/kg) was injected i.p. daily through week 6. **c**, Representative *H&E* images of liver sections from A20 and vehicle treated mice receiving CCl_4_ or oil. **d**, Representative Sirius red images of liver sections from A20 and vehicle treated mice receiving CCl_4_ or oil. **e**, Hydroxyproline assay assessing collagen accumulation in the livers from A20 and vehicle treated mice receiving CCl_4_ or oil (*n* = 5 mice/group, \**p* < 0.05, \*\**p* < 0.01, \*\*\**p* < 0.001 by ANOVA). **f,** Plasma level of ALT from A20 and vehicle treated mice receiving CCl_4_ or oil (*n* = 5 mice/group, \*\*\**p* < 0.001 by ANOVA). **g-h**, Liver mRNA levels of (**g**) fibrotic markers *Col1a1*, *Timp1*, *Ephb2*, *Acta2*, *Pdgfrβ*, *Fn1* and (**h**) inflammatory markers *Ccr2*, *Il-11*, *Cd11b*, *Il-1β*, *Il-33*, *Il-6*, *Tgf-β1* from A20 and vehicle-treated mice receiving CCl_4_ (*n* = 5 mice/group, \**p* < 0.05, \*\**p* < 0.01, \*\*\**p* < 0.001, \*\*\*\**p* < 0.0001 by Student t test). Data are presented as mean ± SEM, ns = not significant.

To determine if A20 could reverse established liver fibrosis in CCl_4_-injured *WT* mice, we administered 20 mg/kg of A20 i.p. once daily starting three weeks after initiating CCl_4_ treatment and continued daily for another three weeks (**Fig. 5b**). H&E stains of liver sections from these mice showed a strong reduction in mononuclear cell accumulation in CCl_4_-injured animals that were dosed with A20 compared to those dosed with vehicle (**Fig. 5c**). Sirius red and measurement of hydroxyproline revealed that collagen accumulation in the livers of A20 treated CCl_4_-injured mice was significantly reduced compared to the vehicle treated counterparts (**Fig. 5d,e**), although plasma ALT was not significantly different between the two groups (**Fig. 5f**). Gene expression analysis showed that mRNA levels of fibrotic (*Col1a1*, *Timp1*, *Ephb2*, *Acta2*, *Pdgfrβ, Fn1)* and inflammatory *(Ccr2*, *Cd11b*, *Il-11*, *Il-1β*, *Il-33*, *Il-6, Tgf-β1*) markers were significantly reduced in the livers from CCl_4_-injured mice treated with A20 compared to the vehicle dosed counterparts (**Fig. 5g,h**). This data indicates dosing animals with an Eph-Ephrin tetramerization inhibitor can effectively reverse established hepatic fibrosis caused by chemical injury.

### A20 mitigates fibrosis in GAN diet-injured livers

We next tested if A20 could reduce MASH hepatic fibrosis caused by a high fat diet. Here, young *WT* mice were either maintained on a normal chow diet or switched to the obesogenic GAN diet for 38 weeks. Beginning on week 28, after MASH fibrosis was established in the GAN fed animals, mice were dosed with either 10 mg/kg of A20 or vehicle only 3 times a week for the remaining 10 weeks (**Fig. 6a**). H&E assessment of liver sections showed that while vehicle-dosed GAN diet fed mice as expected exhibited steatosis with massive hepatocyte ballooning compared to the normal chow fed animals, the livers from GAN fed A20-treated mice appeared much less fattened, consistent with a non-significant trend towards reduced liver-to-body weight ratio in these animals (**Fig. 6b,c**). Sirius red staining and hydroxyproline measurements showed greatly elevated collagen deposition in the vehicle-dosed GAN diet fed mice that was significantly reduced in A20-dosed counterparts (**Fig. 6d,e**). Further, while plasma ALT and AST were not different among the two different GAN fed groups, cholesterol was significantly reduced in the A20 dosed mice compared to vehicle (**Fig. 6f,g**), and expression of fibrotic and inflammatory markers were all significantly down-regulated in the livers from GAN fed, A20 dosed mice (**Fig. 6h,i**). A20 treatment of GAN fed mice also significantly reduced myeloid cell populations normally present in MASH livers (**Fig. 6j**). We conclude dosing mice with A20 in mice fed a MASH-inducing diet will reduce steatosis, inflammation, and fibrosis.

**Fig. 6:**
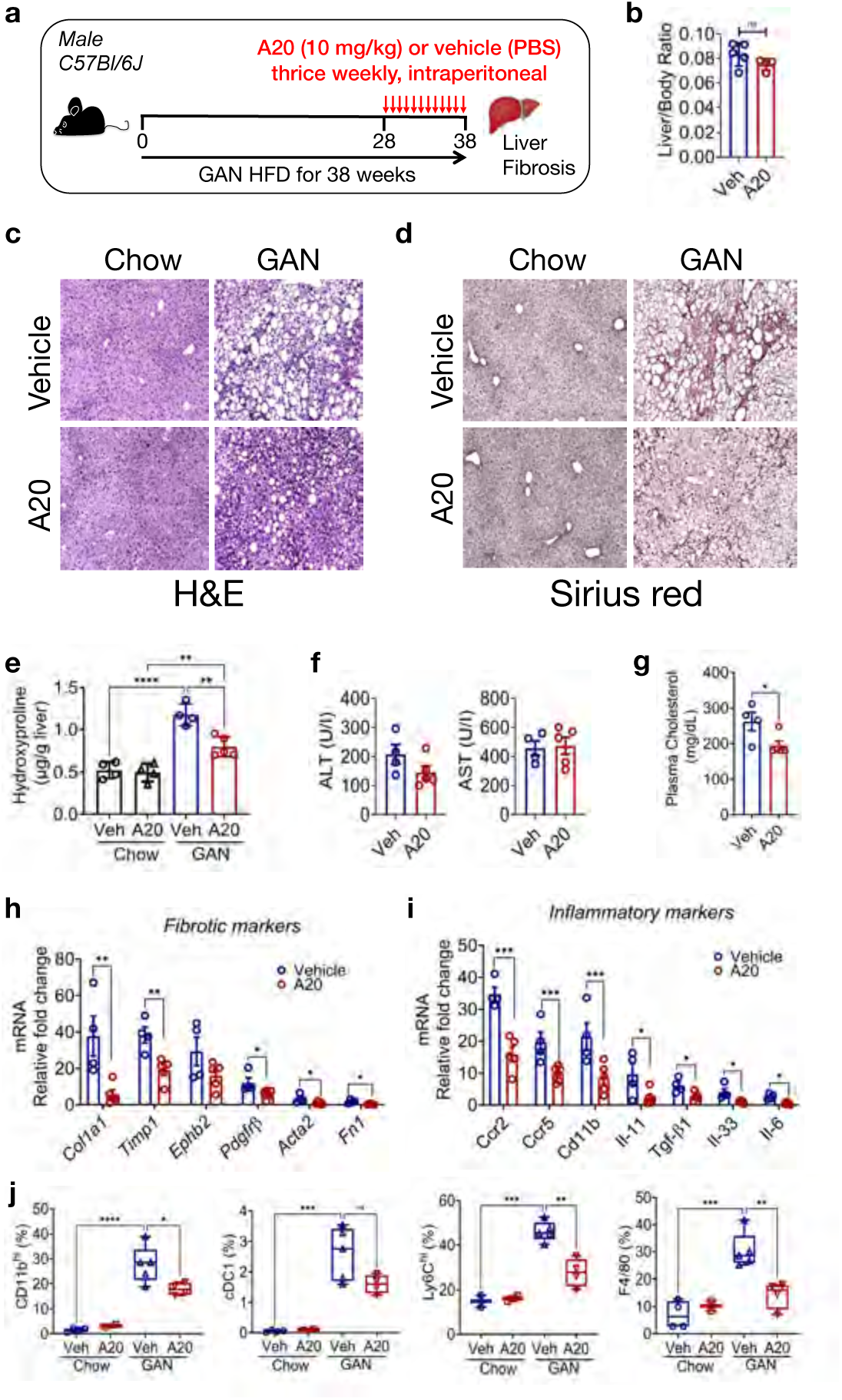
Tetramerization inhibitor A20 mitigates GAN diet-induced MASH fibrosis in mice. **a**, Study outline for GAN HFD diet-induced liver fibrosis model in which male C57BL6/J mice were fed a GAN diet for a total of 38 weeks. Twenty-eight weeks into the GAN diet, either vehicle or A20 tetramerization inhibitor (a 2xHCl salt of A20 was dissolved in PBS and dosed at 10 mg/kg) was injected i.p. 3x weekly for the remaining 10 weeks of the study. **b**, Liver to body weight ratio in vehicle and A20 treated GAN fed mice (n=4-5 per group). **c-d**, Representative images of (**c**) *H&E* stained and (**d**) Sirius red stained liver sections from vehicle and A20 treated mice fed the GAN and chow diets. **e**, Hydroxyproline assay assessing collagen accumulation in the livers of vehicle and A20-treated mice (*n* = 4-5 mice/group, \*\**p* < 0.01, \*\*\*\**p* < 0.0001 by ANOVA). **f-g**, Plasma levels of (**f**) ALT and AST and (**g**) cholesterol from vehicle and A20 treated mice fed the GAN and chow diets (*n* = 4-5 mice/group, \**p* < 0.05, by Student t test). **h-i**, Liver mRNA levels of (**h**) fibrotic markers *Col1a1*, *Timp1*, *Ephb2*, *Pdgfrβ*, *Acta2*, *Fn1* and (**i**) inflammatory markers *Ccr2*, *Ccr5*, *Cd11b*, *Il-11*, *Tgf-β1*, *Il-33*, *Il-6* from vehicle and A20 treated mice fed the GAN diet (*n* =4-5 mice/group, \**p* < 0.05, \*\**p* < 0.01, \*\*\**p* < 0.001 by Student t test). **j**, Immunophenotyping of macrophage subsets (CD11b, Ly6C, F4/80) and dendritic cells (cDC1) from livers of vehicle and A20 treated mice fed the GAN and chow diets (n = 4-5 mice/group, \**p* < 0.05, \*\**p* < 0.01, \*\*\**p* < 0.001, \*\*\*\**p* < 0.0001 by ANOVA). Data are presented as mean ± SEM, ns = not significant.

As GAN diet fed *Ephb2-/- KO* mice are protected from obesity, we noticed that treatment of GAN fed *WT* mice with A20 also significantly reduced body weight compared to the vehicle treated mice without affecting food intake (**Extended Data Fig. 14a,b**). The reduced body weight in A20 dosed GAN fed mice was accompanied by a significant reduction of epididymal (visceral) and inguinal (subcutaneous) white fat deposits compared to vehicle dosed counterparts (**Extended Data Fig. 14c**). Glucose and insulin tolerance tests (GTT and ITT) indicated a non-significant trend for improved glucose and insulin resistance in the A20 dosed mice (**Extended Data Fig. 14d,e**).

Altogether, our data suggest that A20 might, in addition to mitigating hepatic inflammation and fibrosis, also ameliorate metabolic dysfunction commonly observed in animals fed an obesogenic diet.

## Discussion

Fibrosis is associated with poor outcome in patients with MASH, and for this reason the identification of critical signaling pathways involved in fibrogenesis is a priority for the development of potential anti-scaring therapeutics. We have previously showed that the receptor tyrosine kinase EphB2 promotes the development of hepatic, renal, and dermal fibrosis^12,13,18,40^. Here, we report that EphB2 is up-regulated in livers from humans with advanced MASH and in murine models of MASH fibrosis. We dissected the contribution of EphB2 in MASH fibrosis by combining genetic knockout, point mutation, conditional, and knockdown approaches with clinically relevant models by feeding mice either the obesogenic GAN diet for up to 38 weeks or the shorter acting non-obesogenic CDAA diet for 8-12 weeks. We found global deletion of *Ephb2* attenuated GAN diet-induced steatosis, inflammation, and fibrosis likely because of the unexpected lack of obesity observed in these mice, which indicate additional roles for this receptor in metabolism outside of the liver. Nevertheless, as the livers from *Ephb2-/- KO* mice fed the non-obesogenic CDAA diet were protected from inflammation and fibrosis, our data positions EphB2 as a critical hub regulating steatosis, inflammation and fibrosis in MASH and in obesity. The tyrosine kinase catalytic activity of EphB2 is central for this hub-like function in MASH, as kinase-overactive F620D mice presented with excessive hepatic inflammation and fibrosis while the kinase-dead K661R mouse mimicked the knockout and was protected.

Digging deeper into the critical role for EphB2 tyrosine kinase forward signaling in MASH, we found that deleting receptor expression in the hepatocyte compartment did not affect diet-induced liver fibrosis, which is in contrast to prior studies^11^. This discrepancy could be due to different diets employed in the two studies and/or because we undertook a genetic conditional strategy with a widely used Albumin-Cre driver to delete *Ephb2* in hepatocytes, while the other group used an AAV RNA targeting approach. Notwithstanding, we noticed that EPHB2 expression was mostly elevated in HSCs in advanced MASH patients and its expression in these cells becomes strongly up-regulated by TGF-β/SMAD signaling, which was validated by SMAD complex binding to the putative promoter region of human *EPHB2*. Moreover, we found siRNA silencing of *EPHB2* in HSCs blocked TGF-β-mediated HSC-to-myofibroblast transdifferentiation *in vitro*. This is likely due to a crosstalk between Eph/Ephrin signaling and TGFβ/SMAD signaling as co-immunoprecipitation and proximity ligation assay experiments confirmed a physical EphB2-TGFβR2 protein-protein association in hepatic stellate cells.

Furthermore, knocking down *EPHB2* attenuated the ability of TGF-β to induce SMAD activation in human HSCs/myofibroblasts, and revealed that the top down-regulated genes in TGF-β stimulated *EPHB2* deleted HSCs were mostly involved in fibrogenesis, including *EPHB2* itself. Overall, our mechanistic studies suggest a model whereby the EPHB2 receptor interacts with TGFβR2, and together by interacting with their respective Ephrin and TGF-β ligands work together to facilitate canonical TGFβ/SMAD signaling to drive stellate cell activation and transdifferentiation into profibrotic myofibroblasts.

Finally, for the sake of developing effective anti-fibrotic therapies to combat MASH, we demonstrate novel Eph-Ephrin tetramerization inhibitor A20, which is orally available^22^, shows potent ability to counter fibrosis in either chemical-induced or diet-induced liver injury models. We anticipate further development and testing of Eph-Ephrin tetramerization inhibitors will lead to promising new methods to treat MASH fibrosis as well as obesity and perhaps other metabolic and fibroinflammatory disorders.

## Materials and Methods

### Human samples

De-identified human livers samples from patients with MASLD/MASH at various stage of fibrosis (F0 – F4, modified Kleiner-Brunt staging methodology) 41, reviewed by a liver pathologist, were obtained from the University of Utah Pathology Archives. The protocol was approved by the University of Utah Institutional Review Board and determined to be exempt from human subjects’ research (IRB # 00117459).

### Mice

Animal experiments were performed in compliance with guidelines of Indiana University Institution Animal Care and Use Committee (IACUC) protocol number 24131 MD/R/HZ/ E/EX and University of Utah IACUC protocol number 1688. Male and female C57BL/6J mice [RRID:IMSR_JAX:000664], aged 8-12 weeks, were bred in-house or purchased from Jackson laboratory (Bar Harbor, ME). *Ephb2-/-* and littermate controls *(Ephb2+/+)* mice on the C57BL6/J background has been described previously 42 and bred in-house. *Ephb2*-kinase overactive mice (*Ephb2^F620D/F620D^*) and *Ephb2*-kinase dead (*Ephb2^K661R/K661R^*) originally in the CD1/129 background 27,43, were backcrossed for 10 generations into C57BL6/J strain. To generate hepatic stellate cells/mesenchymal stromal cells *Ephb2* knockout mice, *Ephb2^loxP/loxP^* mice were crossed with the tamoxifen-inducible Cre recombinase driven by mouse promoter *Col1a2Cre*^ERT2^ (RRID:IMSR_JAX :029235) or *PdgfrβCre^ERT2^* (RRID:IMSR_JAX :029684). Hepatocyte-specific *Ephb2* knockout mice were generated by crossing *Ephb2^loxP/loxP^* mice with Alb-Cre mice (RRID:IMSR_JAX :003574). All the *Cre* mouse lines used in this study were purchased from Jackson Laboratories (Bar Harbor, ME). Littermates with floxed alleles but without Cre were used as wild type controls. Where appropriate, both Cre inducible mouse line and controls animals were administered tamoxifen 75mg/kg for 5-7 consecutive days and hepatic fibrosis induced a week later. All animals were housed in a specific pathogen-free environment, with no more than five animals per cage under controlled light (12h light and 12h dark cycle), temperature (24 ± 2°C), humidity (50 ± 10%) conditions, and provided water and food *ad libitum* throughout all experiments.

### Diets and chemical induction of MASH fibrosis

Eight to ten weeks old male or female mice were fed a normal chow diet, the obesogenic Gubra-Amylin NASH (GAN) diet (D09100310, Research diets, Inc)) with additional fructose (23.1g/L D-fructose (Sigma-Aldrich, F0127) and glucose (18.9g/L D-glucose (Sigma-Aldrich, G8270) supplemented in drinking water for 22 to 26 weeks or the Choline Deficient L-Amino Acid defined high-fat diet (CDAA-HFD) (A16092003, Research Diets, Inc) for 8 to 12 weeks. In some experiments, mice were subjected to chemical induction of liver fibrosis using carbon tetrachloride (CCl_4_) intraperitoneal injection (Sigma-Aldrich 1:10 v/v diluted in olive oil at 2µl/g body weight) three times a week for 4 weeks. Control mice were injected with olive oil, and all animals were sacrificed 48 hours after the final injection of CCl_4_. At the end of each experiment, animals were sacrificed by isoflurane (Fluriso, USP) inhalation, blood and livers were harvested for subsequent analysis.

### Immunofluorescence analysis of human MASH liver

Paraffin-embedded sections of human liver were deparaffinized, rehydrated, and treated for 20 min with boiling citrate antigen retrieval buffer (Sigma-Aldrich). Slides were blocked with 5% donkey serum in 1x Tris-buffered-saline containing 0.1% Triton-X-100 (TBS-T) for 1h at room temperature and incubated overnight at 4 °C with goat anti-EphB2 (AF467, R&D System), rabbit anti-phosphoEphB1/B2^Y594 + Y596^ (Abcam), and mouse anti-αSMA (1:200 dilution in 1% BSA TBS-T) antibodies. Slides were washed and incubated for 1 h with donkey anti-mouse IgG NorthernLights-493, donkey anti-rabbit or donkey anti-goat IgG Northernlghts-557 (1:500 dilution in 1% BSA-TBST). Slides were washed three times with 1xTBS-T and mounted with VECTASHIELD Antifade Mounting Medium with DAPI (VectorLabs). Images were acquired using a Nikon A1R confocal microscope and analyzed using the ImageJ software (NIH).

### Immunohistochemistry

Immunohistochemistry staining was performed in 6-µm thick tissue sections. After deparaffinization and rehydration in ethanol solutions, antigen retrieval was performed in citrate buffer solution (pH 6.0) for 30 minutes at 95°C. Endogenous peroxidase activity was blocked using hydrogen peroxide solution at room temperature, followed by washing in phosphate-buffered solution saline (PBS) and passage through distilled water. The slides were incubated overnight with rabbit anti-phosphoEphB1/B2^Y594 + Y596^ antibody diluted (1:50) in 5% BSA-TBST. Slides were washed three time in TBS-T and incubated with the secondary antibody EnvisionTM Flex/HRP (with Streptavidin Peroxidase Solution, Agilent, DAKO) at 37°C for 1 hour. Slides were stained with 3, 30-diaminobenzidine tetrahydrochloride (DAB-Agilent, DAKO), counterstained with Harris hematoxylin (Thermo Scientific™), and mounted with a resinous mounting medium (Permount™) dissolved in xylene. High power images were acquired using an EVOS M5000 wide-field microscope (ThermoFisher).

### Single cell preparation of liver tissue for flow cytometry

After mice were euthanized using isoflurane, blood was collected by cardiac puncture, the abdomen was exposed and the liver collected, rinse with PBS and weighed. Liver was subsequently transferred in approximately 3ml of serum-free RPMI-1640 containing Collagenase D (10mg/ml; Sigma) and DNase (1mg/ml; Sigma) and incubated in a rocking platform for 45 min at 37°C. The liver extract was mashed through a 70µm filter, the cell re-suspended in RPMI-1640 containing 10% FBS and centrifuged at 1600 rpm for 5 min. The supernatant was discarded, and the pellet re-suspended in approximately 4 ml of 70% Percoll, then transferred in 15 ml conical tube, carefully overlay with 4 ml of 30% Percoll and centrifuged 1600 rpm for 25 min with the brake turned off. The non-parenchymal cells suspension from the Percoll interface were removed and mixed with 10 mL of RPMI-1640 containing 10% FBS and the cells were centrifuged at 1600 rpm for 5 min. Red blood cells (RBC) were removed from the pelleted single cell suspensions of livers non-parenchymal cells by incubation in an ammonium chloride-based 1x RBC lysis buffer (Thermofisher, eBioscience). The cells were again pelleted and mixed with FACS buffer (2% BSA, 2mM EDTA in PBS), then stained with Zombie-NIR viability dye (BioLegend) per manufacturer’s instructions to discriminate live vs dead cells. To prevent non-specific Fc binding, the cells were incubated with Fc Block (anti-mouse CD16/32, clone 93, Biolegend) for 15 min followed by the indicated antibodies cocktail for 60 min in the dark on ice: CD45 (FITC, clone S18009F, Biolegend), CD11b (BVC421, clone M1/70, Biolegend), F4/80 (APC, clone BM8, Biolegend), TIM4 (PerCP/Cy5.5, clone RMT4-54, Biolegend), Ly6C (PE, clone HK1.4, Biolegend), MHCII (BV605, clone M5/114.15.2, Biolegend), CD11c (BV785, clone N418, Biolegend) and Ly6G (PE/Cy7, clone 1A8, Biolegend). After surface staining, cells were fixed with a paraformaldehyde-based fixation buffer (BioLegend). Flow cytometric acquisition was performed on a BD Fortessa X20 flow cytometer (BD Biosciences) and data analyzed using FlowJo software (Version 10.8.1; Tree Star Inc).

### Histology

Livers specimens were fixed in 10% buffered formalin and multiple paraffin-embedded 6µm sections were stained with *H&E* Masson’s Trichrome and Sirius red according to standard protocols. Images were taken at ten randomly selected field using an EVOS M5000 widefield microscope and analyzed using the ImageJ software (NIH).

### Hydroxyproline measurement

Hepatic collagen content was measured using the hydroxyproline assay as previously described^44^.

### Mouse plasma collection and analysis

Blood obtained by cardiac puncture was transferred in Eppendorf tubes containing 50µl of heparin, centrifuged at room temperature for 30 min at 10000 rpm and plasma collected from the supernatant fraction. Plasma samples were snap frozen and stored at −80°C until analysis. Plasma samples were processed in a single batch for determination of alanine aminotransferase (ALT), aspartate aminotransferase (AST) cholesterol and triglyceride levels using a DC Element chemistry analyser (HESKA).

### Primary human hepatic Stellate cells

Primary human hepatic stellate cells (HSCs) were purchased from ScienCell Research Laboratories (Carlsbad, CA, USA, catalog number 5300) and cultured in a specialty medium (ScienCell #530). LX2 cell lines were purchased from Sigma-Aldrich and cultured in the fibroblast growth kit-low serum (ATCC, VA, catalog number PCS-201-041) constituted with the following ingredients: Fibroblast basal medium, ascorbic acid, fetal bovine serum, rhFGF-basic, L-glutamine, and hydrocortisone. These cells were maintained at 37°C in a humidified 5% CO_2_ incubator and medium were changed every 48 hours. HSCs between passage 1 and 3 were used in this study.

### TGF-β stimulation of HSCs

HSCs were plated in 24-well tissue culture plates and cultured as described above until at least 70% confluency was reached. Subsequently, HSCs were serum-starved overnight in Dulbecco’s Modified Eagle Medium/F12 (DMEM/F12) containing 0.5% Bovine Serum Albumin (BSA) and exposed to recombinant human TGF-β1, TGF-β2, and TGF-β3 all at 10 ng/ml (R&D Systems) for 3h, 6h, 12h, 24h, and 48h. At each time point, vehicle and TGF-β stimulated HSCs were used for assessment of *EPHB2* and *ACTA2*/αSMA mRNA and proteins levels by RT-qPCR or western blot.

### Inhibition of canonical TGF-β/SMAD pathway

To assess the effect of TGF-β/SMAD canonical pathway on EphB2 expression, HSCs were incubated with the TGF-β type I receptor (TβRI) inhibitors SD208 (1µM), SB525334 (200nM), and the SMAD3 inhibitor SIS3 (5µM) (all from Tocris) for 4h before stimulation with recombinant human TGF-β1 (10ng/ml) for 48h. RNA was extracted to assess *EPHB2* mRNA expression by RT-qPCR.

### siRNA transfections of HSCs

HSCs (70% confluency) were transfected with *EPHB2,* SMAD2, and *SMAD3* Mission®-esiRNA (Sigma Aldrich, USA) delivered into the cells using the lipofectamine^®^ RNAiMAX (Invitrogen, USA) reagent according to the manufacturer’s instruction. Mission®-esiRNA complexes targeting eGFP were used as non-targeting control. After 48H incubation, transfected HSCs were stimulated with recombinant human TGF-β1 at 10 ng/ml for another 48H. Cells were used for immunofluorescence staining or harvested for RNA and proteins extractions.

### RNA extraction, cDNA synthesis and qPCR

Total RNA was extracted using RNA Stat60^®^ (Tel-Test Inc, USA) following standard phenol-chloroform procedure, and further purified using the NucleoSpin RNA-II purification kit (Machery-Nagel) following the manufacturer’s instructions.

Complementary DNAs were generated by reverse transcription using Superscript reverse transcriptase-II (Invitrogen, USA) as previously described 13. Real-time qPCR reactions were performed using Quantitect SYBR Green PCR reagent (Qiagen) and qPCR reaction plates run in the Applied BioSystems FAST 7000 Sequence detection system (ABI Prism FAST 7000). Primer sequences are shown in Supplementary Table 1 and 2, and were designed using the web-based application PrimerBank 45. Transcripts were normalized to different housekeeping genes, *glyceraldehyde-3-phosphate dehydrogenase* (GAPDH) and *hypoxanthine-guanine phosphoribosyl transferase* (*HPRT*). Messenger RNA expression levels were calculated using the 2^-ΔΔCt^ method.

### *EPHB2* locus-specific Chromatin Immunoprecipitation

HSCs were serum-starved overnight and then treated with rhTGF-β1 at 5 ng/ml for 3 hr. Control cells were not treated. Cells were then harvested for the ChIP assay using the ChIP-IT High Sensitivity kit (Cat 53040, Active Motif) as per the manufacturer’s recommendation. Briefly, proteins were cross-linked to DNA in living cells with formaldehyde, cells were lysed, homogenized with a chilled Dounce homogenizer to release nuclei. The nuclear pellet was resuspended in sonication buffer and shared on ice using a digital sonicator (Branson SFX 150, EMERSON) at amplitude setting of 40% with pulse rate 30 seconds on and 30 seconds off to yield DNA fragment sizes of 200 - 1200 bp. The shared chromatin (∼30 µg) samples were immunoprecipitated using 4 µg of SMAD3 antibodies (Abcam, ab28379) or purified anti-human IgG Fc (Cat 410701, BioLegend) overnight at 4°C. The immune complexes were collected by incubation with Protein G Agarose beads (Cat 37499, Active Motif), followed by reverse crosslinking, digestion with Proteinase K and DNA purification. Enrichment of *EPHB2* promoter sequences in the immunoprecipitated eluted DNA was determined by qPCR using the set of primers designed based on the estimated binding region of EPHB2 (Table S1).

### Immunofluorescence of HSCs

HSCs controls and HSCs-transfected with *siEPHB2* were seeded on chamber slides, treated with vehicle or TGF-β1 as described above and fixed with 4% PFA for 10–15 min. The cells were then washed three times with TBS-T, blocked for one hour in blocking solution (5% donkey serum; 0.1% triton-X-100 in TBS) and incubated overnight at 4 °C with a primary antibody against EPHB2 (AF467, R&D System), and α-SMA (clone 1A4, R&D system; 1:200 in blocking solution). The next day, samples were washed three times with TBS-T incubated for one hour with donkey anti-goat IgG NorthernLights-493 and donkey anti-mouse IgG, NorthernLights-557 (1:500) (R&D System) in blocking solution. Samples were washed three times with TBS-T and mounted with VECTASHIELD Antifade Mounting Medium with DAPI (Vectorlabs). Images were acquired with a Nikon A1R confocal microscope.

### RNA sequencing

RNA sequencing was performed as previously described (Egal et al 2014) Total RNA from EphB2-knockout and control HSCs stimulated with rhTGF-β1 were extracted as described above. RNA sequencing was performed by Novogene on an Illumina NovaSeq platform using a paired-end 150-bp sequencing strategy. The human GRCh38 and mouse GRCm38 genome and gene annotation files were downloaded from Ensembl release 98 and a reference database was created using STAR version 2.7.2c with splice junctions optimized for 150 base pair reads 46. Optical duplicates were removed from the paired end FASTQ files using clumpify v38.34 and reads were trimmed of adapters using cutadapt 1.16 47. The trimmed reads were aligned to the reference database using STAR in two pass mode to output a BAM file sorted by coordinates. Mapped reads were assigned to annotated genes using featureCounts version 1.6.3 48. The output files from cutadapt, FastQC, FastQ Screen, Picard CollectRnaSeqMetrics, STAR and featureCounts were summarized using MultiQC to check for any sample outliers 49. Differentially expressed genes were identified using a 5% false discovery rate with DESeq2 version 1.26.0 50. Pathways were analyzed using the fast gene set enrichment package and Ingenuity Pathway Analysis 51 (Qiagen, Inc). The RNA-seq datasets have been deposited in the GEO database (GSE274897, GSE274898, GSE274899).

### Immunoprecipitation and immunoblotting

Samples were lysed in radioimmunoprecipitation assay (RIPA) buffer supplemented with protease and phosphatase inhibitors (Thermo Scientific). Protein concentrations were determined using Pierce Rapid Gold BCA protein Assay Kit (Thermo Scientific), diluted to a concentration of 2,000µg/ml of protein. EphB2 proteins were immunoprecipitated using the Dynabeads Protein A Immunoprecipitation kit (Invitrogen) with 2µg of anti-EphB2 antibody (R&D Systems, AF467), heated for 15 mins at 70°C and cooled down on ice. Subsequently, proteins were separated by SDS–PAGE electrophoresis and transferred to 0.2µm PVDF membranes (Millipore, Merck). Membranes were incubated in blocking solution (1% gelatin in 1X TBS with 0.5% Tween 20, Sigma) for 1.5 hour, and primary antibody was incubated overnight at 4°C in the blocking solution. The antibodies and their concentrations are the following: anti-EPHB2 (R&D Systems, AF467, 1:2,000), and anti-TGFβR2 (E5M6F, 1:1000, CST-41896, Cell Signaling Technology) Following several washes in TBST (TBS with 0.5% Tween 20), horseradish peroxidase (HRP)-labeled secondary antibodies (1:5,000) were incubated for 1 h at room temperature in 10% nonfat dry milk TBST. Membranes were developed with the Super Signal West Pico PLUS Chemiluminescent western-blotting substrate (Thermo Scientific).

Cells were lysed in ice-cold RIPA buffer containing protease and phosphatase inhibitor cocktails (Thermo Fisher Scientific). Protein concentration in cell lysates were determined using a Pierce^TM^ rapid gold bicinchoninic acid (BCA) protein assay kit from ThermoFisher Scientific. Proteins were loaded and resolved on a 10% Tris-Glycine SDS-PAGE gels, electro-transferred onto nitrocellulose membrane and incubated overnight at 4°C with specific antibodies. These antibodies and their concentrations are the following: anti-EPHB2 (R&D Systems, AF467, 1:2,000), anti-α-SMA (D4K9N, 1:10,000), anti-phospho-SMAD2/3 (D27F4, 1:10,000) anti-SMAD2/3, (D7G7, 1:1000), anti-phospho-AKT (D9E, 1:1000), anti-AKT (40D4, 1:10,000) anti-SMURF2 (D8B8, 1:1000) and anti-vinculin (E1E9V; 1:10,000) all from Cell Signaling Technology. Immunoreactivity was revealed by incubation with host-specific secondary antibodies conjugated with horseradish peroxidase, followed by detection with SuperSignal West Pico Chemiluminescent Substrate from ThermoFisher. Images were acquired using an Amersham imager 680 Chemi system from GE Healthcare.

### Kinome array and analysis

PamGene’s KinomePro approach measure the ability of active kinases in a cell lysate sample to phosphorylate specific peptides imprinted on PamChip® microarray 52. The PamChip® consists of 4 identical arrays, each array containing 144 (STK) or 196 (PTK) phosphosites. HSCs (70% confluency) were transfected with *EPHB2* Mission®-esiRNA (Sigma Aldrich, USA) delivered into the cells using the lipofectamine^®^ RNAiMAX (Invitrogen, USA) reagent according to the manufacturer’s instruction. Mission®-esiRNA complexes targeting eGFP were used as non-targeting control. After 48H incubation, transfected HSCs were further stimulated with recombinant human TGF-β1 at 10 ng/ml for 48H. Whole-cell lysates were prepared according to PamGene instructions.

PamChip PTK and STK arrays were performed by PamGene. Measurements of kinome activity were performed on a PamStation-12 by PamGene 53. Briefly, the PamChip PTK array was processed in a single-step reaction in which 5.0 microgram of protein lysate was dispensed onto PTK array dissolved in protein kinase buffer (PamGene proprietary information) and additives including 1% BSA, 10 mmol/L dithiothreitol, FITC conjugated pY20 antibody, and 400 mmol/L ATP. The STK array was processed in a two-step reaction in which 1.5 microgram of protein lysate was used with protein kinase buffer (PamGene proprietary information) supplied with 1% BSA, primary STK antibody mix, and 400 mmol/L ATP (sample mix). After an initial incubation of 110 minutes, the reaction mix was removed, and the secondary FITC-labeled antibody mix was added. In both arrays, software-based image analysis (BioNavigator software V.6.3 from PamGene) integrates the signals obtained within the time course of the incubation of the kinase lysate on the chip into one single value for each peptide for each sample (exposure time scaling). Peptide phosphorylation kinetics (for PTK) and its variations (for PTK/ STK) were used to remove low signal peptides as quality control analysis (QC). Individual peptide phosphorylation intensities were log2 transformed. The peptides with significant differences in phosphorylation intensity (HSCs-*EPHB2*-silenced vs HSCs, using test, P < 0.05) are visualized as either volcano and bar plots or as heatmap using the R heatmap package.

Upstream kinase analysis (UKA) of PTK and STK data was done using default setting of the PamApp (PTK or STK UKA 2018 V.4.0) on BioNavigator Analysis software tool as described before 54. The analysis is based on documented kinase–substrate relationships (from iviv database and literature-based protein modifications such as HPRD, PhosphoELM, PhosphositePLUS, Reactome, UNIPROT) complemented with “in silico” predictions that are retrieved from the phosphoNET database. The analysis generated three major parameters calculated by the PamApp: (i) Median kinase statistic (MKS) depicts the overall change of the peptide set that represents a kinase. For instance, a larger positive value indicates a larger activity in treated cells compared with controls. (ii) Mean significance score (MSiS) indicating the significance of the change represented by the mean kinase statistic (s) between two groups (using 500 permutations across sample labels). (iii) Mean specificity score (MSpS) indicates the specificity of the mean kinase statistics with respect to the number of peptides used for predicting the corresponding kinase (using 500 permutations across target peptides).

The final ranking of the kinases was based on Median Final Score (MFS), which was calculated by addition of MSiS and MSpS. Top predicted kinases from significant (MFS>1.3) PTK and STK peptide sets are represented on phylogenetic tree of the human protein kinase family generated by Coral (http://phanstiel-lab.med.unc.edu/ CORAL/; ref. 30).

### Migration assay

We devised a custom void filling migration assay like the migration assays provided by Ibidi (GmbH). A rectangular silicone membrane (1mm thickness, 5 mm x 15 mm) was placed in a 6-well plate, and primary human HSCs were plated around the silicone membrane at cell density approximately 50% confluence. HSCs (80% confluency) were transfected with *EPHB2* Mission®-esiRNA (Sigma Aldrich, USA) delivered into the cells using the lipofectamine^®^ RNAiMAX (Invitrogen, USA) according to the manufacturer’s instruction. Mission®-esiRNA complexes targeting eGFP were used as non-targeting control. After 48 hours upon confirming 100% confluence, silicone membrane was removed, and void was created (Time 0 hour). Cell migration was monitored every 6 hours, and images were taken at the consistent locations by referencing a marking line etched at the edge of silicone membrane. Average migratory distance was calculated from 5 different locations.

### TGF-β/SMAD reporter assay

The SBE reporter kits (79806 and 79578, BPS Biosciences) were used to monitor the activity of the TGF-β/SMAD signaling pathway in human HSCs, following the manufacturer’s instruction. HSCs were seeded into a 24-well plate, cultured to 70- 80% confluency and transfected with *EPHB2* Mission®-esiRNA (Sigma Aldrich, USA) delivered into the cells using the lipofectamine^®^ RNAiMAX (Invitrogen, USA) according to the manufacturer’s instruction. Mission®-esiRNA complexes targeting eGFP were used as non-targeting control. After 48 hr, transfected HSCs were transduced with SBE luciferase reporter lentivirus and negative control luciferase lentivirus for another 24H. Next, HSCs were stimulated with recombinant human TGF-β1 at 10 ng/ml for 3H and the activation of the TGF-β/SMAD signaling pathway was monitored by measuring luciferase activity using the ONE-GLO luciferase assay system (E6110, Promega) and a luminometer (SpectraMAX iD3 plate reader), following the manufacturer’s instruction. The luciferase activity from the SBE reporter was calculated by subtracting the luminescence from the negative control reporter as a background.

### *In situ* proximity ligation assay

Cells were fixed with 4% paraformaldehyde (Biolegend)) for 15 min and permeabilized for 10 min at room temperature with 0.1% Triton-X 100 (Sigma). The following primaries antibodies were incubated overnight at 4°C: anti-EphB2 (Goat antibody AF467, R&D Systems) and anti-TGFβR2 (Rabbit antibody, clone E5M6F, CST-41896, Cell Signaling Technology). *In situ* proximity ligation assays were performed using the Duolink In Situ reagents (Sigma). Incubation of antibodies, ligation of oligodeoxynucleotides and amplification were performed according to the manufacturer’s instructions. Nuclei were labelled with DAPI and images acquired with a Nikon A1R confocal microscope.

### snRNA-seq data processing

All count matrices and metadata for each sample are publicly available in the Gene Expression Omnibus (www.ncbi.nlm.nih.gov/geo/) under data accession no. GSE212837 10. Briefly, Cellranger count pipeline was applied to the FASTQs to perform alignment against modified transcriptomes based on the mm10 reference builds for mice that contain introns to increase mapping efficiency, and the number of genes detected. Filtered feature-barcode matrices from Cell Ranger were subsequently run through a standard Seurat pipeline for QC. Different QC parameters were used for snRNA-seq data generated using Chromium 3′ Gene Expression V2 Kits using Seurat. For snRNA-seq generated from mouse samples, we included nuclei that fit the following criteria: (i) total number of expressed genes greater than 300 and less than 6000, (ii) <5% of which annotated as mitochondrial, and (iii) total counts greater than 500 and less than 15000. Feature plots were generated using default parameters in Seurat.

### Statistics

Statistical analysis and plotting were done using GraphPad Prism software 9.0.2. Comparisons between more than 2 groups were performed using Kruskal-Wallis One-Way ANOVA and the non-parametric Mann Whitney-U test was used for comparison between 2 non-related groups. Number of biological replicates are indicated in Fig. legends. Data were presented as Mean ± SEM for experiments, and *p* values less than 0.05 were considered significant.

## Acknowledgments

We appreciate the University of Utah and Indiana University cores facilities for assistance with histology, RNA sequencing, confocal microscopy, metabolic phenotyping and flow cytometry studies. We thank Quinian Johanson for excellent animal husbandry, Erika Egal for assistance with histology, and Scott Friedman for providing reagents and resources.

## Author Contributions

P.N.M. conceived the project with input from M.H. Experiments were designed by P.N.M and M.Y., and conducted by M.Y., S.D.K., H.W., A.K., F.S., E.M.K, E.A., S.P., K.J.E., M.N.H., M.G., J.S., H.C.R., S.B., and P.M.M. Early work, A20 tetramer inhibitor, animal models, and various collections by H.W., A.K., F.S., and M.H. Results were analyzed by P.N.M., M.Y., S.D.K., M.G., J.S., H.C.R., S.B., M.K. Figures were designed by P.N.M. Analysis and visualization of single nucleus RNA sequencing data was performed by S.W. The manuscript was written by P.N.M. and M.H., with assistance from M.N.H., H.W., and A.K.

## Lead contact

Further information and requests for resources and reagents should be directed to Lead Contact, Patrice N. Mimche.

## Extended Data Figures

**Extended Data Fig. 1:**
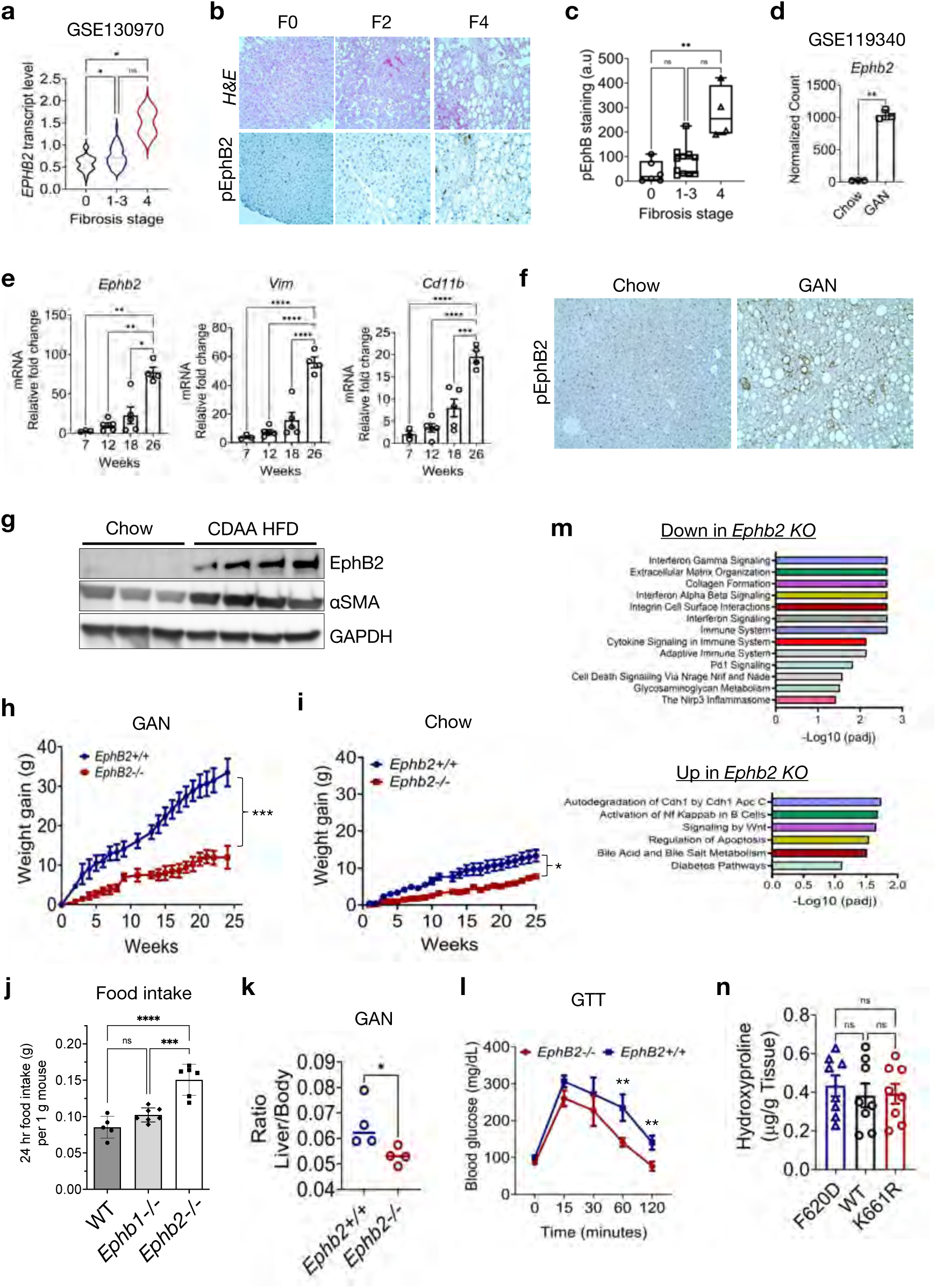
Up-regulated EphB2 expression and activity in human and murine MASH fibrosis and effect of *Ephb2*-/- knockout (*KO*). **a**, Analysis of human MASLD/MASH RNA-seq data (GSE130970) showing that *EPHB2* transcript is elevated in advanced MASH (\**p*<0.05 by ANOVA). **b**, Representative images of liver sections from human with MASLD/MASH at various stage of fibrosis, stained with H&E and phospho-EphB2. Images were taken using a 20x objective. **c**, Quantification of phospho-EphB2 in liver specimen of human with MASLD/MASH (n = 4-10 mice/group, \*\**p* < 0.01 by ANOVA). **d**, Analysis of murine MASH RNA-seq data (GSE119340) showing upregulation of liver *Ephb2* transcript in MASH (n=3/group, \**p*<0.05 by Student’s t-test). **e**, Liver mRNA levels of *Ephb2*, *Vim*, *Cd11b* in mice fed the obesogenic GAN diet at week 7, 12, 18, and 26 (n = 3-5 mice/group, \**p* < 0.05, \*\**p*< 0.01, \*\*\**p*<0.001, \*\*\*\**p*<0.0001 by ANOVA). **f**, Representative images of immunohistochemistry of phospho-EphB2 in liver sections of chow and GAN fed mice at week 26. Images were taken using a 10x objective. **g**, Immunoblot analysis to show protein levels of EphB2, αSMA and GAPDH in liver samples of chow and CDAA diet fed mice for 12 weeks (n=3 mice/group). **h-l**, Body weight gained in *Ephb2+/+* wild-type (*WT*) and *Ephb2-/-* knockout (*KO*) mice fed (**h**) the obesogenic GAN diet for 26 weeks or (**i**) a normal rodent chow for 26 weeks, with (**j**) food intake of normal chow fed mice, (**k**) liver to body weight ratio of GAN fed mice, (**l**) Glucose tolerance test of GAN fed mice at week 22 (n = 4-6 mice/group, \**p* < 0.05, \*\**p*< 0.01, \*\*\**p*<0.001, \*\*\*\**p*<0.0001 by Student’s t-test or ANOVA). **m,** Gene ontology analysis of the top significantly down-regulated or up-regulated pathways in RNA sequencing data from GAN diet fed *Ephb2*+/+ *WT* and *Ephb2*-/- *KO* mice. **n**, Level of hydroxyproline in the livers of *Ephb2-F620D*, *WT*, and *Ephb2-K661R* mice fed the GAN diet for 26 weeks (n = 8 mice/group). Data are presented as mean ± SEM, ns = not significant.

**Extended Data Fig. 2:**
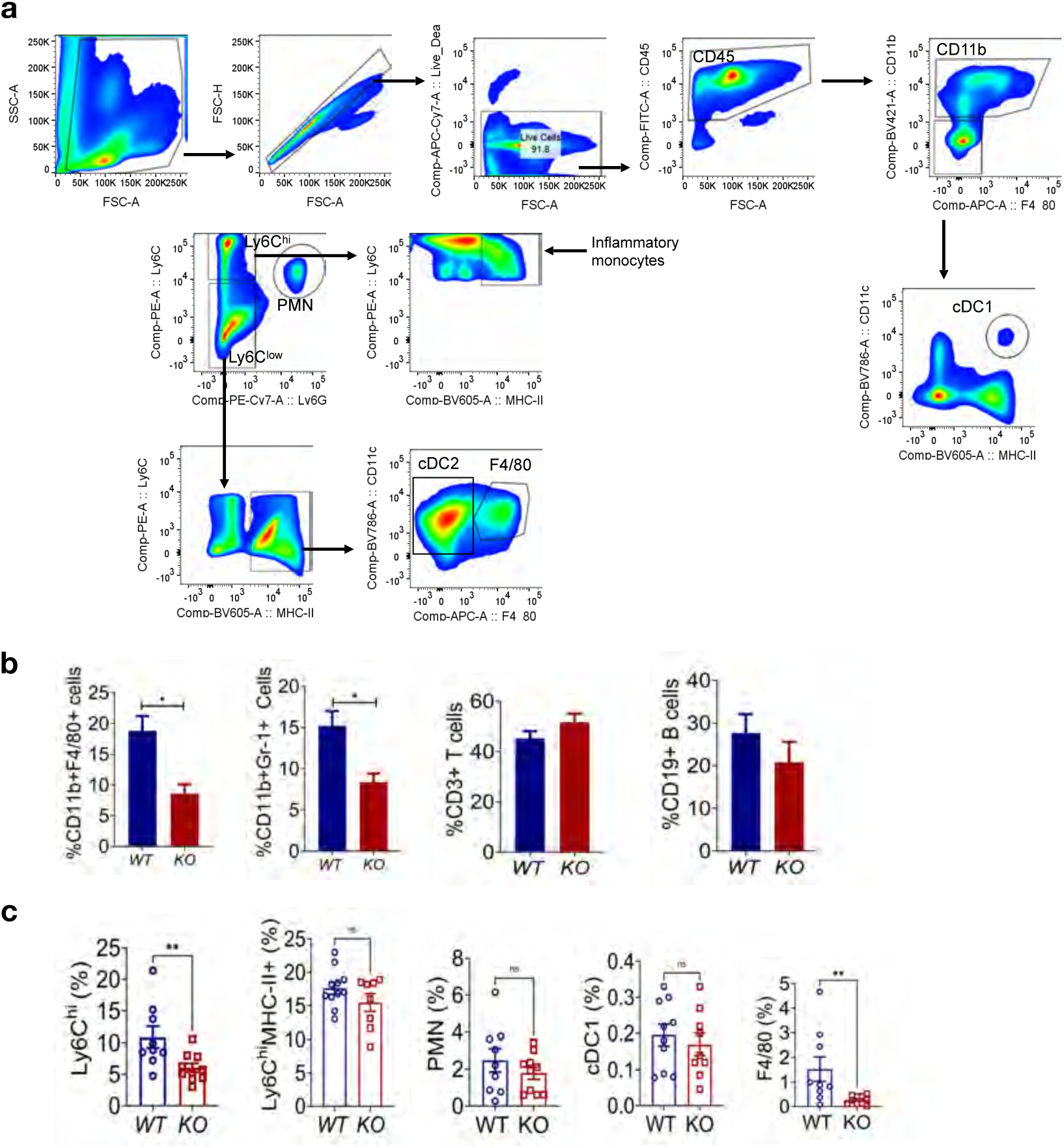
Liver immunophenotyping of myeloid cells in *Ephb2+/+* (*WT*) and *Ephb2*-/- (*KO*) mice fed the GAN and CDAA HFDs. **a**, Gating strategy used to identify various immune cells present in the livers of *WT* and *KO* mice. **b**, Quantification of hepatic myeloid cells subsets and T (CD3) and B (CD19) cells in the livers of *WT* and *KO* mice fed the obesogenic GAN diet for 26 weeks (n = 5 mice/group, \**p*<0.05 by Student’s t-test). **c**, Immunophenotyping of macrophage subsets (Ly6C, PMN-neutrophils, F4/80) and dendritic cells (cDC1) in livers from *WT* and *KO* mice fed the non-obesogenic CDAA diet for 12 weeks (n = 9-10 mice/group, \*\**p*<0.01 by Student’s t-test). Data are presented as mean ± SEM, ns = not significant.

**Extended Data Fig. 3:**
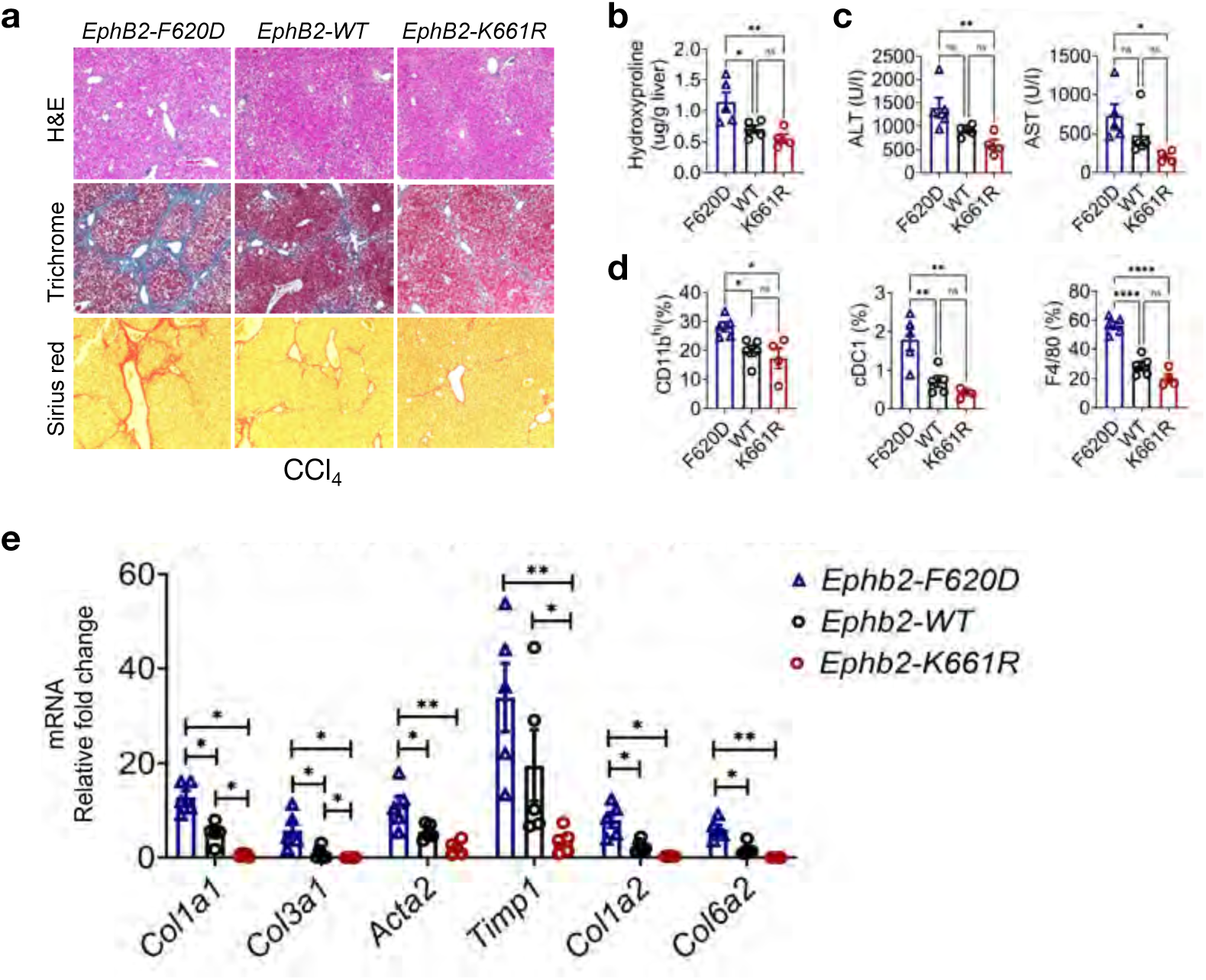
EphB2 tyrosine kinase signaling in CCl_4_ injury-induced fibrosis. **a**, Representative images of liver sections from *Ephb2-F620D* (kinase-overactive point mutant), *Ephb2-WT*, and *Ephb2-K661R* (kinase-dead point mutant) mice that received CCl_4_ for 4 weeks and stained with *H&E*, Masson’s Trichrome and Sirius red. **b-d**, Levels of (**b**) liver hydroxyproline, (**c**) plasma liver damage enzymes ALT and AST, and (**d**) liver myeloid cell population subsets (CD11b^hi^, cDC1, F4/80) from *F620D*, *WT*, and *K661R* mice that received CCl_4_ for 4 weeks (n = 4-5 mice/group, \**p*<0.05, \*\**p*<0.01, \*\*\*\**p*<0.0001 by ANOVA). **e**, Liver mRNA levels of fibrotic markers *Col1a1*, *Col3a1*, *Acta2*, *Timp1*, *Col1a2*, *Col6a2* in CCl_4_-injured *F620D*, *WT*, and *K661R* mice (*n* = 4-5 mice/group, \**p*<0.05, \*\**p*<0.01 by ANOVA). Data are presented as mean ± SEM, ns = not significant.

**Extended Data Fig. 4:**
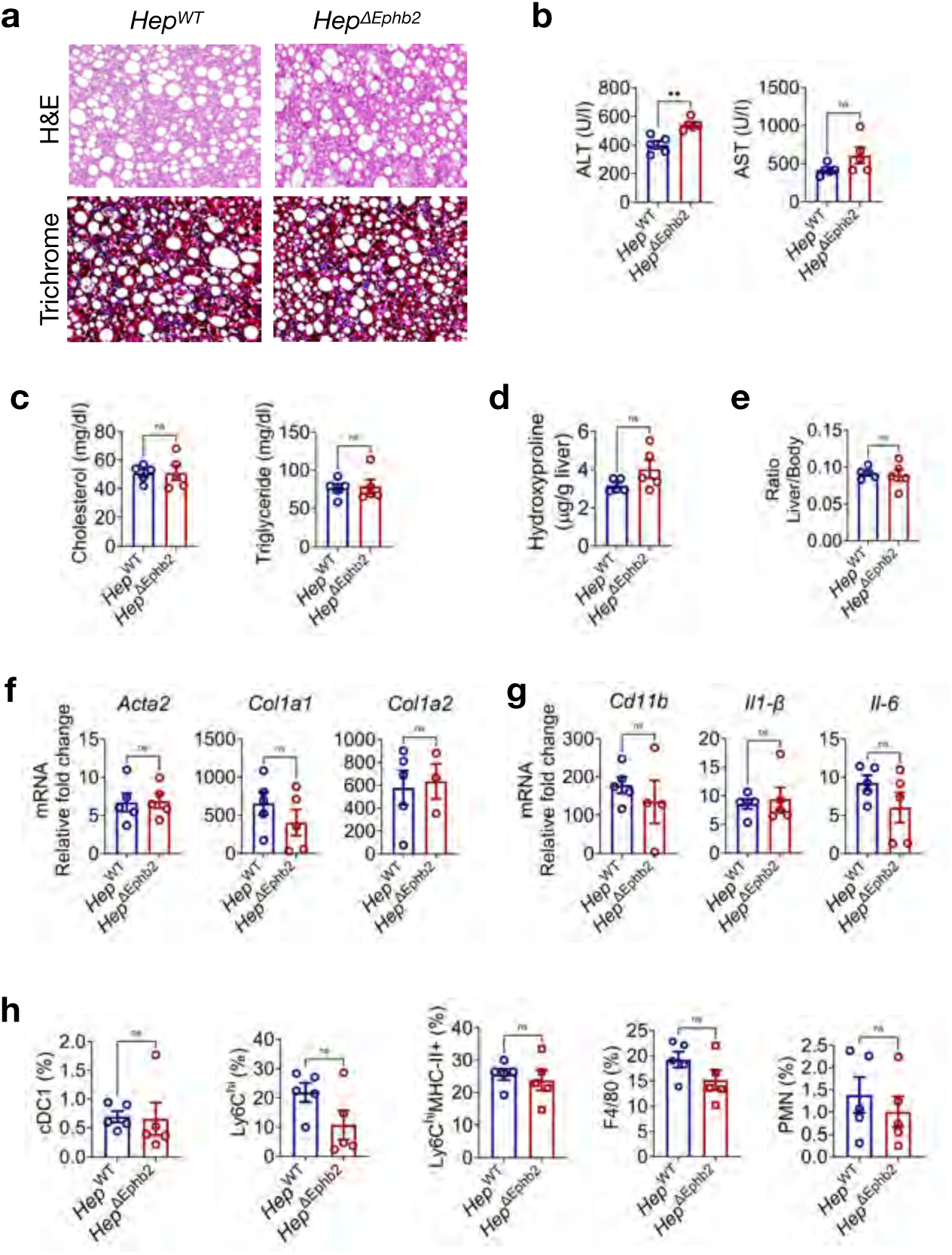
Deletion of *Ephb2* in hepatocytes does not mitigate CDAA diet-induced MASH in mice. **a**, Representative images of liver sections from *Hep^WT^* and *Hep^ΔEphb2^* mice fed the CDAA HFD for 12 weeks stained with *H&E* and Masson’s Trichrome to assess tissue damage and collagen deposition. **b-c**, Plasma levels of (**b**) liver damage enzymes ALT and AST and (**c**) cholesterol and triglycerides from the CDAA HFD fed *Hep^WT^* and *Hep^ΔEphb2^* mice (n = 4-7 mice/group, \*\**p*< 0.01 by Student’s t-test). **d-g**, Liver (**d**) hydroxyproline levels, (**e**) body weight ratio, (**f**) mRNA levels of fibrotic markers *Col1a1*, *Col1a2*, *Acta2*, and (**g**) mRNA levels of inflammatory markers *Cd11b*, *Il-1β*, *Il-6* from CDAA HFD fed *Hep^WT^* and *Hep^ΔEphb2^* mice (n = 4-6 mice/group). **h**, Immunophenotyping of various myeloid cells subsets in the livers of CDAA fed *Hep^WT^* and *Hep^ΔEphb2^* mice (n = 4-7 mice/group). Data are presented as mean ± SEM, ns = not significant.

**Extended Data Fig. 5:**
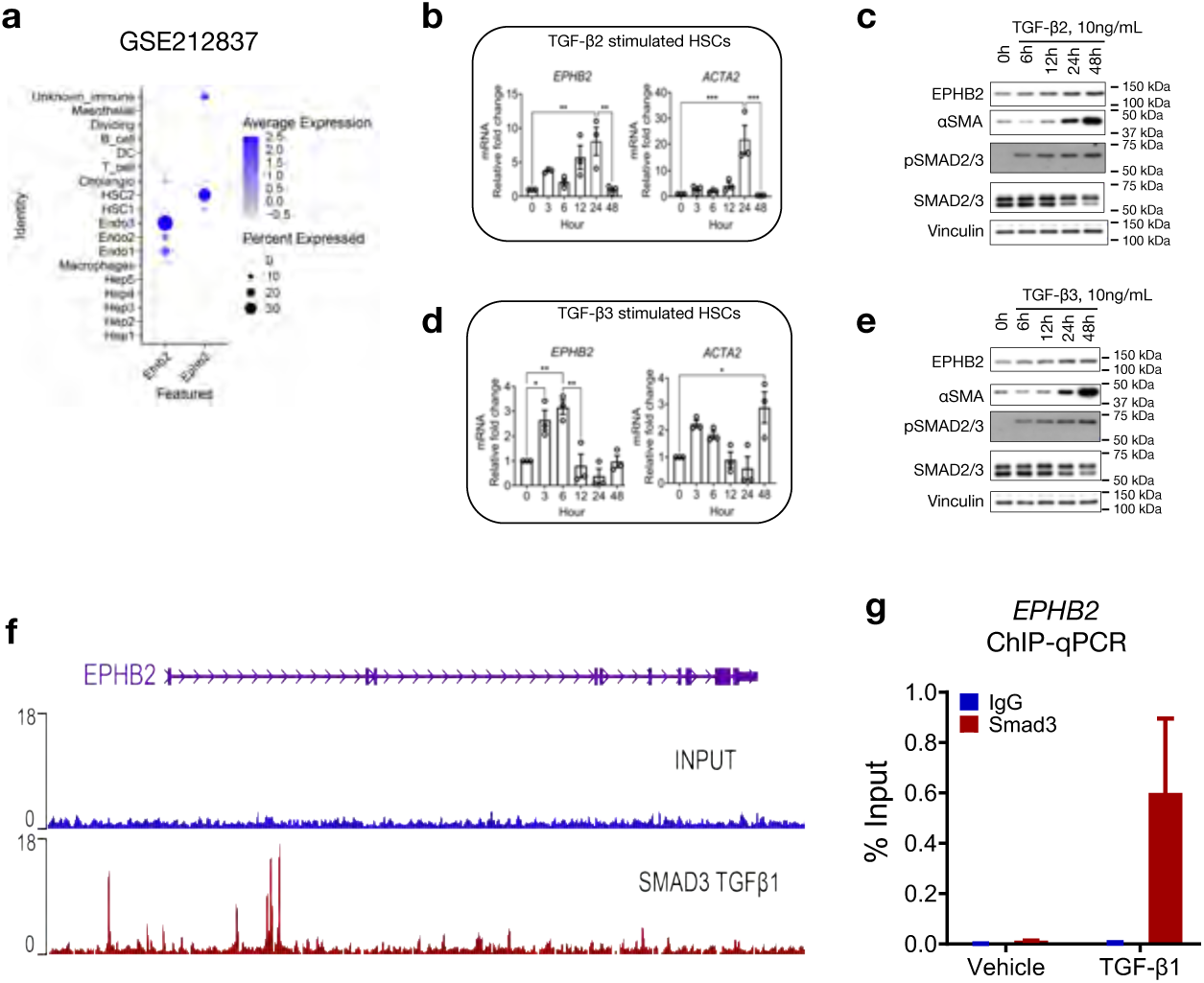
Expression of Ephb2 in hepatic cell types is up-regulated by TGF-β2 and TGF-β3 and consistent with SMAD3 binding sites within the EPHB2 promotor region. **a**, Dot plot showing expression of EphB2 and one of its ligands Ephrinb2 (*Efnb2*) across different hepatic cell types in mouse snRNA-seq analyzed from published dataset GSE212837 of mice fed a western diet, with average expression levels normalized to those from livers of standard chow diet fed mice. b-c, (**b**) Messenger RNA levels of *EPHB2* and *ACTA2* and (**c**) protein levels of EPHB2, αSMA, pSMAD2/3, SMAD2/3, and VINCULIN in TGF-β2 (10 ng/ml) stimulated primary human HSCs at various time points (n = 3, \*\**p*<0.01, \*\*\**p*<0.001 by ANOVA). Data is presented as mean ± SEM. **d-e**, (**d**) Messenger RNA levels of *EPHB2* and *ACTA2* and (**e**) protein levels of EPHB2, αSMA, pSMAD2/3, SMAD2/3, and VINCULIN in in TGF-β3 (10 ng/ml) stimulated primary human HSCs at various time points (n = 3, \**p*<0.05, \*\**p*<0.01 by ANOVA). Data is presented as mean ± SEM. **f**, Accessibility peaks for the transcription factor SMAD3 on the promoter of human *EPHB2* locus identified from a publicly available ChIP-seq dataset GSE38103 of TGF-β1 stimulated LX2 cells. **g**, Enrichment of SMAD3 binding sites on the promoter of human *EPHB2* validated by ChIP-qPCR in TGF-β1 stimulated primary human HSCs.

**Extended Data Fig. 6:**
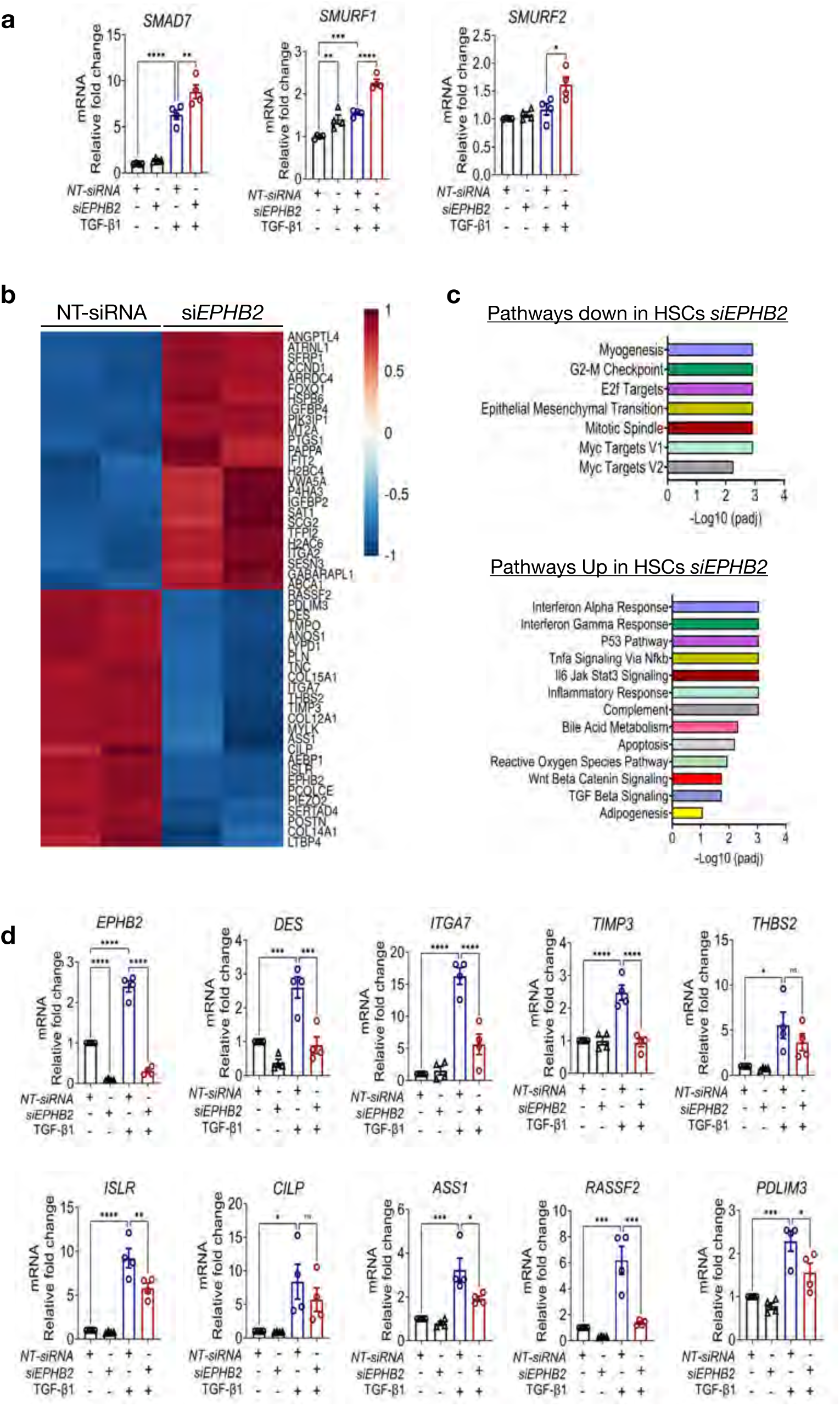
Effects of EPHB2 silencing in TGFβ1 stimulated primary human HSCs. **a**, Messenger RNA levels of *SMAD7*, *SMURF1*, and *SMURF2* measured in primary human HSCs treated with Mission® *EPHB2*-siRNA (SIGMA) and then stimulated with TGF-β1 at 10 ng/ml, with data normalized to HSCs treated with non-targeting Mission® NT-siRNA (n = 4/group, \**p* < 0.05, \*\**p*< 0.01, \*\*\**p*<0.001, \*\*\*\**p*<0.0001 by ANOVA). **b**, Heatmap depicting the top 40 differentially expressed genes in bulk RNA sequencing of TGF-β1 stimulated HSCs lines treated with either non-targeting control NT-esiRNA or experimental *EPHB2*-esiRNA. **c**, Gene ontology analysis of the top significantly down-regulated or up-regulated pathways in RNA sequencing data in **b. d**, Quantitative PCR validation of the top fibrogenic genes up-regulated in the RNA sequencing data set of TGF-β1-stimulated non-targeting control HSCs compared to TGF-β1-stimulated *EPHB2*-silenced HSCs and the unstimulated controls (n = 4/group, \**p*<0.05, \*\**p*<0.01, \*\*\**p*<0.001, \*\*\*\**p*<0.0001 by ANOVA). Data is presented as mean ± SEM.

**Extended Data Fig. 7:**
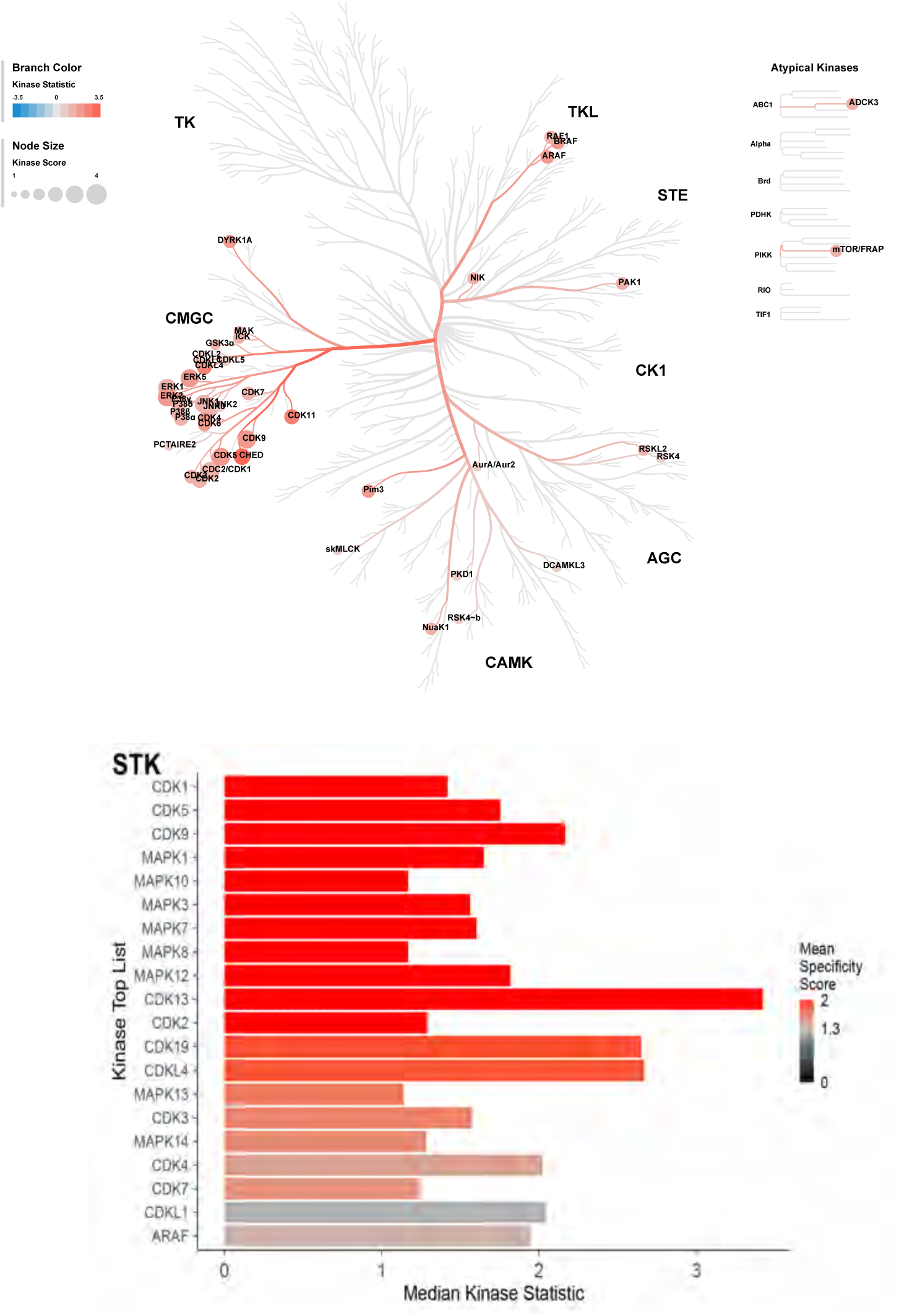
Kinome mapping to identify kinase networks differentially regulated in TGF-β1 stimulated control and *EPHB2* silenced human HSCs. Paralogous relationships between differentially expressed kinases in TGF-β1 stimulated non-targeting control and *EPHB2*-silenced primary human HSCs stimulated with 10 ng/ ml TGF-β1 for 48 hr. CDKs and MAPKs families of serine/threonine kinases were significantly affected by knocking down *EPHB2* in TGF-β1 stimulated HSCs.

**Extended Data Fig. 8:**
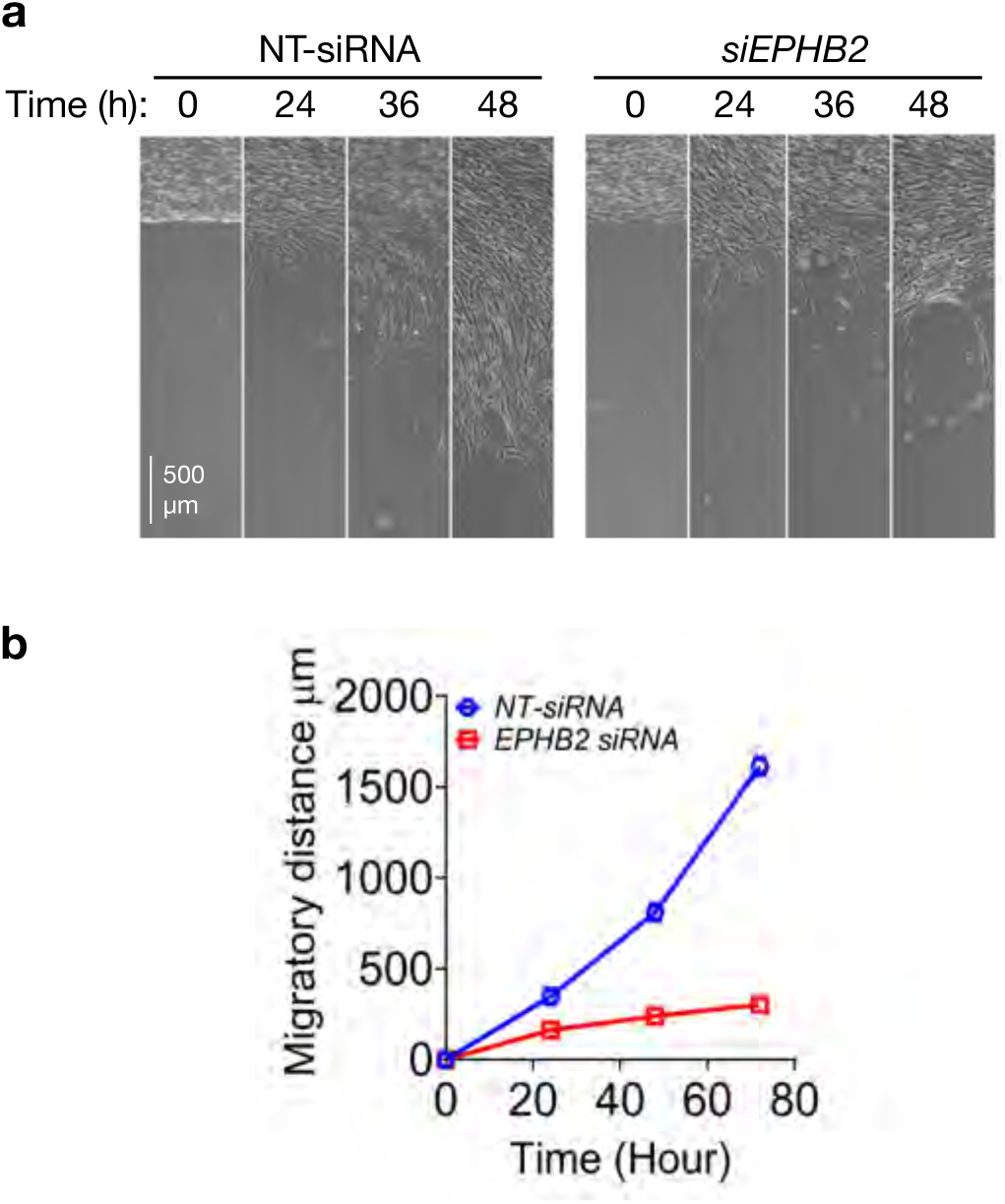
Cell migration is reduced in *EPHB2* silenced human HSCs. **a**, Live cell imaging of HSCs migration in non-targeted control NT-siRNA and *EPHB2*-siRNA transfected cells at various time points following creation of a void to entice cell movement. **b**, Graph of the data (n = 3/group, \**p*<0.05, \*\**p*<0.01 by Student’s t-test). Data are presented as mean.

**Extended Data Fig. 9:**
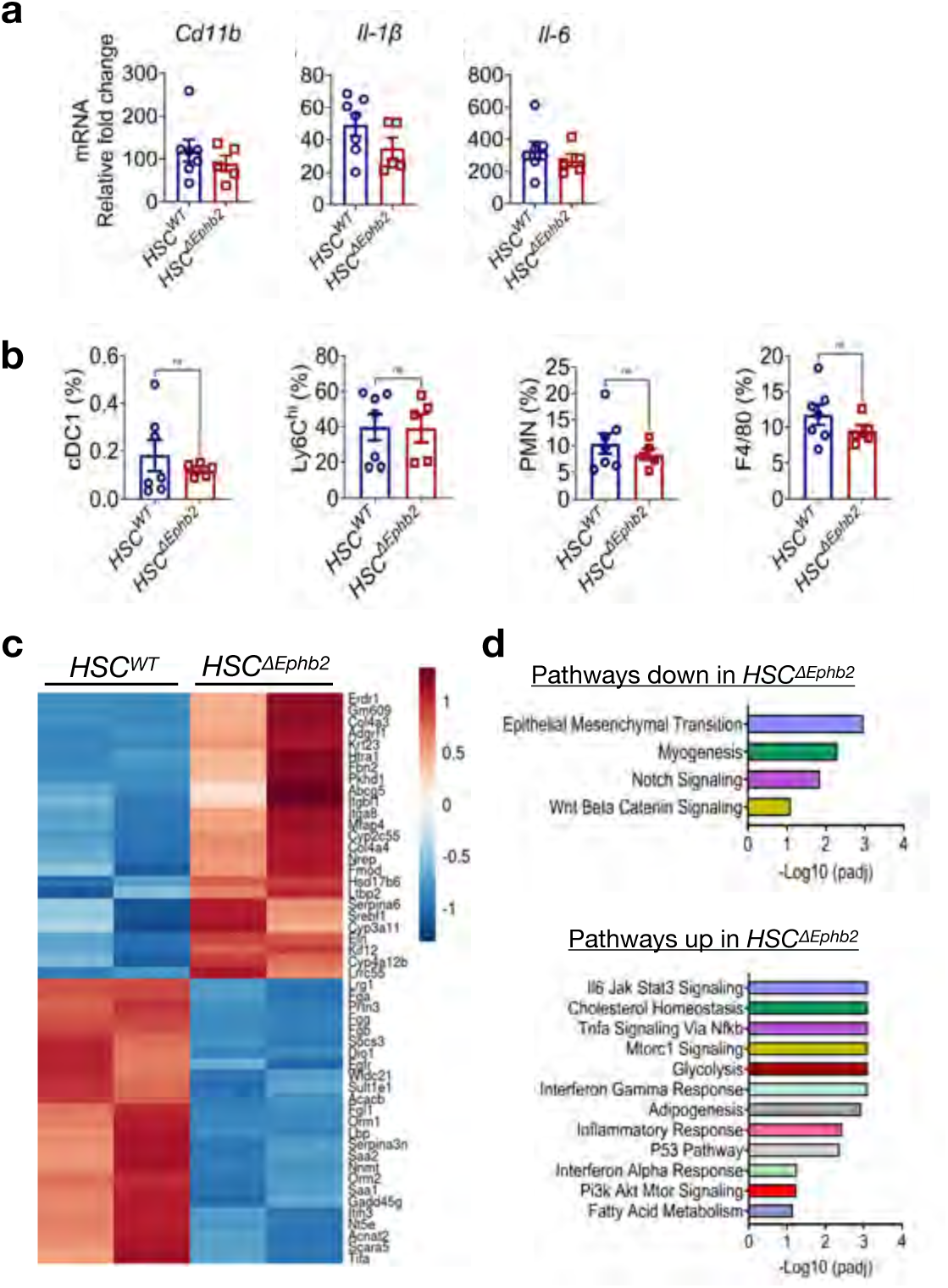
Deletion of *Ephb2* in hepatic stellate cells mitigates CDAA diet-induced MASH fibrosis. **a**, Liver mRNA levels of *Cd11b, Il-1β, and Il-6* in Cre-negative *Ephb2*^loxP/loxP^ (*HSC^WT^*) *WT* control mice and Cre-positive *HSC^ΔEphb2^* mice were injected i.p with tamoxifen and then fed the CDAA diet for 8 weeks (n = 5-7 mice/group). **b**, Percentage of myeloid cell populations in the livers from *HSC^WT^* and *HSC^ΔEphb2^* mice fed the CDAA diet (n = 5-7 mice/group). Data are presented as mean ± SEM. ns = not significant. **c**, Heatmap depicting the top 40 differentially expressed genes from bulk RNA sequencing of livers tissues of *HSC^WT^* and *HSC^ΔEphb2^* mice fed the CDAA diet. **d**, Gene ontology analysis of the top significantly down-regulated or up-regulated pathways in RNA sequencing data in **c.**

**Extended Data Fig. 10:**
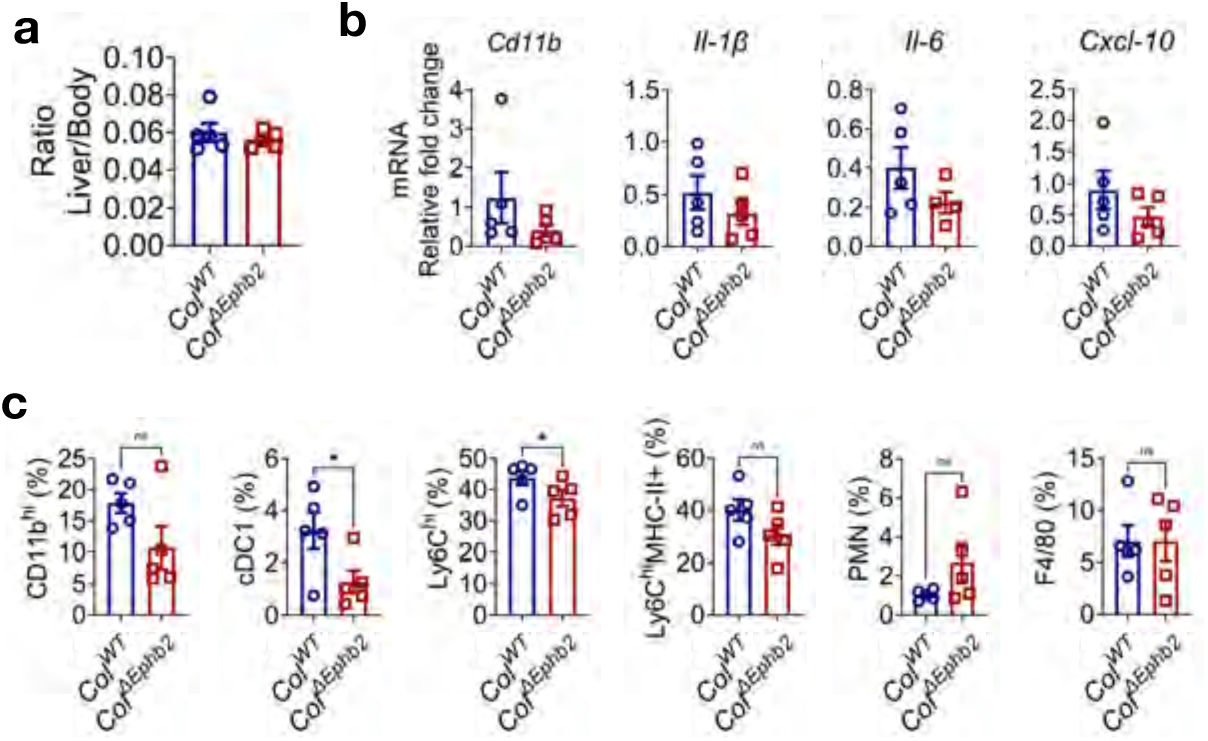
Deletion of *Ephb2* in hepatic stellate cells mitigates GAN diet-induced MASH fibrosis. **a**, Liver to body weight ratio of livers from Cre-negative *Ephb2*^loxP/loxP^ (*Col^WT^*) *WT* control mice and Cre-positive *Col^ΔEphb2^* mice were injected i.p with tamoxifen and then fed the GAN diet for 22 weeks. **b**, Liver mRNA levels of *Cd11b*, *Il-1β*, *IL-6*, *Cxcl-10* in *Col^WT^* and *Col^ΔEphb2^* mice fed the GAN diet. **c**, Immunophenotyping of myeloid subsets (CD11b, Ly6C, MHC-II, F4/80), dendritic cells (cDC1), and neutrophils (PMN) in livers from *Col^WT^* and *Col^ΔEphb2^* mice fed the GAN diet (n = 5 mice/ group, \**p*<0.05 by Student’s t-test). Data is presented as mean ± SEM, ns = not significant.

**Extended Data Fig. 11:**
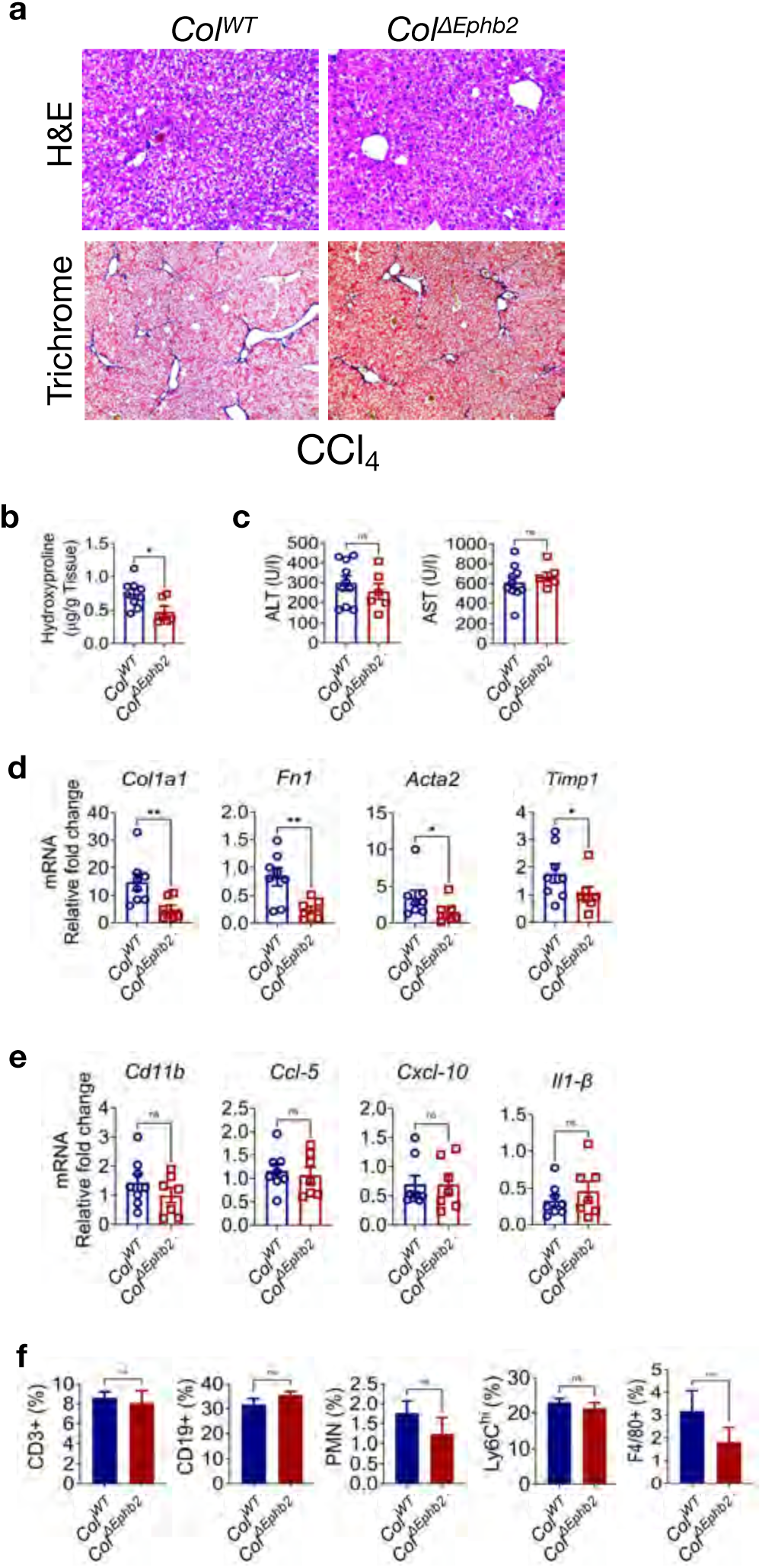
Deletion of *Ephb2* in HSCs attenuates CCl_4_-induced hepatic fibrosis. Cre-negative *Ephb2*^loxP/loxP^ (*Col^WT^*) *WT* control mice and Cre-positive *Col^ΔEphb2^* mice were injected i.p with tamoxifen and then mice treated with CCl_4_ for 4 weeks. **a**, Representative images of liver sections from *Col^WT^* and *Col^ΔEphb2^* mice stained with *H&E* and Masson’s Trichrome to assess tissue damage and collagen deposition. **b**, Hydroxyproline quantified in the livers of *Col^WT^* and *Col^ΔEphb2^* mice. **c**, Plasma levels of ALT and AST in *Col^WT^* and *Col^ΔEphb2^* mice. **d-e**, Liver mRNA levels of (**d**) fibrotic markers *Col1a1*, *Fn*, *Acta2*, *Timp1,* and (**e**) inflammatory markers *Cd11b*, *Ccl-5*, *Cxcl-10*, *Il-1β* from *Col^WT^* and *Col^ΔEphb2^* mice. **f**, Immunophenotyping of T cell (CD3), B cells (CD19), and macrophages subset (Ly6C, F4/80, PMN,) in the livers of Cre- and Cre+ mice receiving (n = 6-10 mice/group, \**p*<0.05, \*\**p*<0.01 by Student’s t-test). Data is presented as mean ± SEM, ns = not significant.

**Extended Data Fig. 12:**
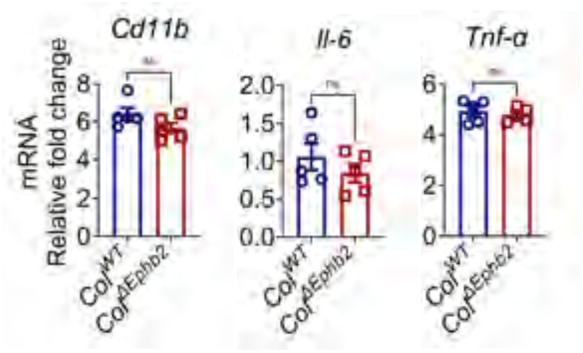
Deletion of *Ephb2* in liver mesenchymal cells when fibrosis is already established attenuates CDAA HFD diet-induced MASH fibrosis. Liver mRNA levels of *Cd11b, IL-6 and Tnf-α* in *Col^WT^* and *Col^ΔEphb2^* mice fed the CDAA diet for 12 weeks and injected i.p with tamoxifen during week 8 (n = 5 mice/group). Data are presented as mean ± SEM, ns = not significant.

**Extended Data Fig. 13:**
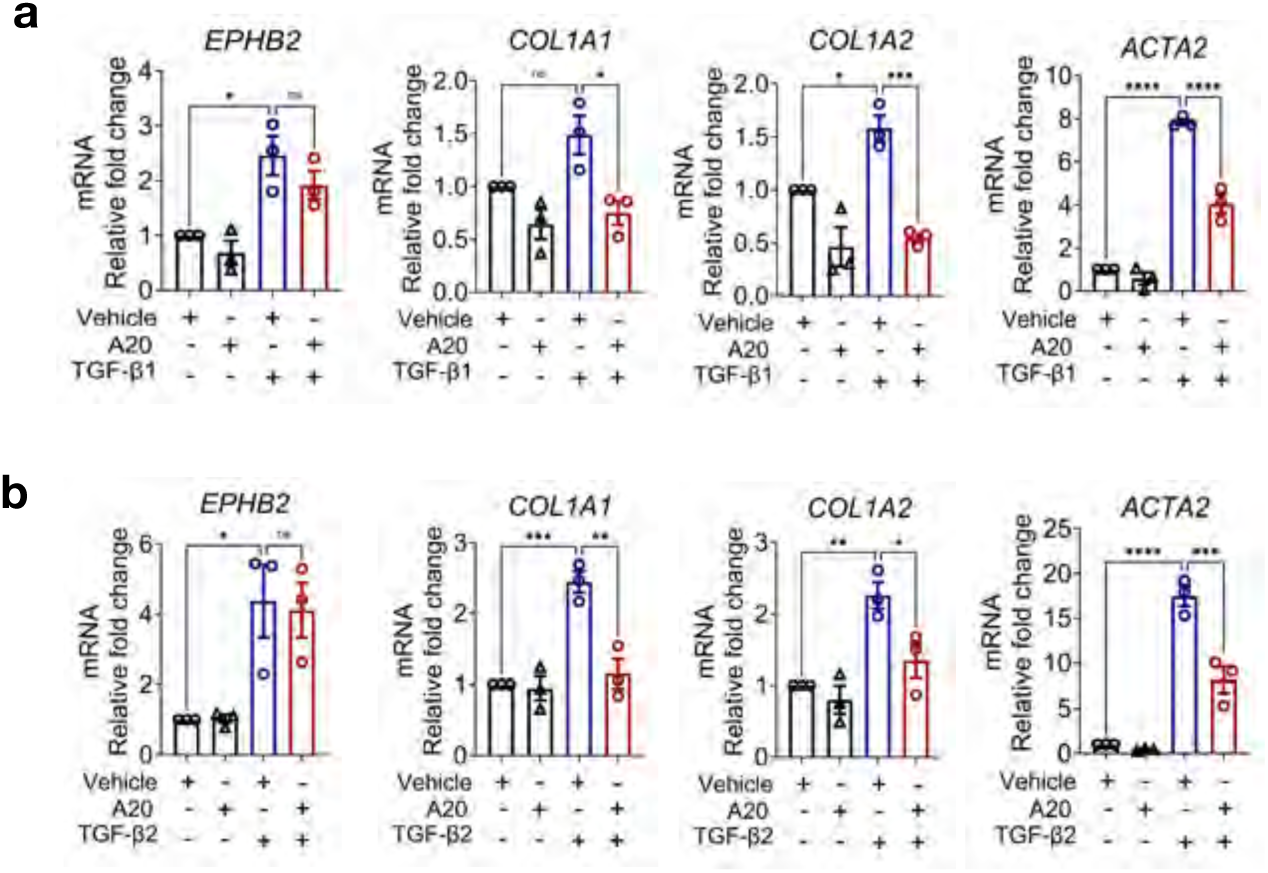
A20 abrogates TGF-β-induced human HSCs activation. Human hepatic stellate cells were first stimulated with 10 ng/ml TGF-β1 (**a**) or TGF-β2 (**b**) for 48 hr and then further incubated in the presence or absence of 5 µM of A20 tetramerization inhibitor compound for an additional 24 hr. RNA was extracted and subjected to quantitative PCR to assess gene expression of fibrotic markers *EPHB2*, *COL1A1*, *COL1A2*, *ACTA2* (n = 3/group \**p* < 0.05, \*\**p*< 0.01, \*\*\**p*<0.001, \*\*\*\**p*<0.0001 by ANOVA). Data represent Mean ± SEM.

**Extended Data Fig. 14:**
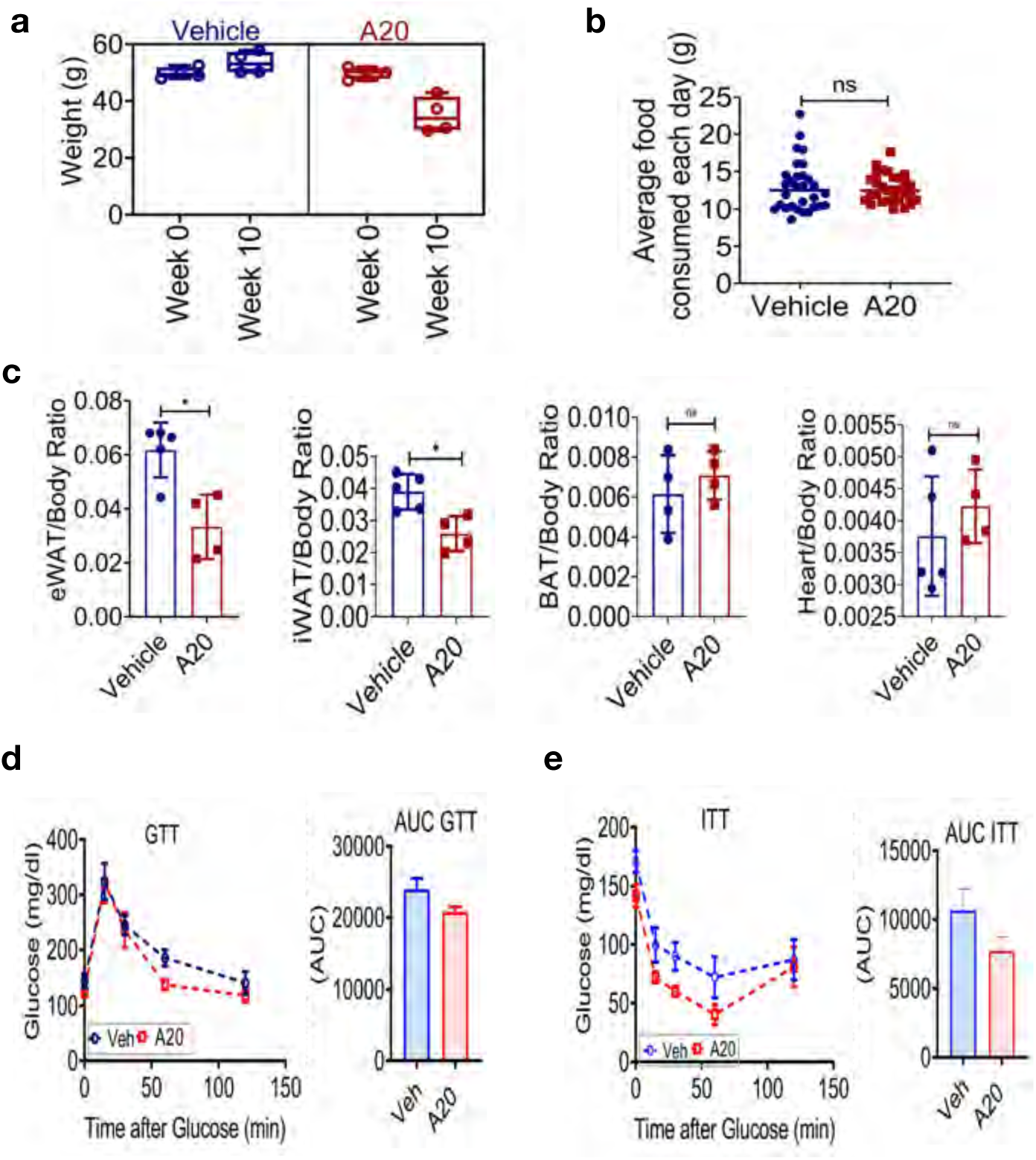
A20 reduces fat depot and improves glucose homeostasis in mice fed the obesogenic GAN diet. Male C57BL6/J were fed the obesogenic GAN diet for a total of 38 weeks. Treatment regimen consists of A20 (10 mg/kg) injection i.p. (3x weekly) from week 28 to week 38. **a**, Weight was recorded at week 28 before treatment with vehicle or A20 (dosing week 0), and then again 10 weeks later after all thirty dosings of vehicle or A20 were administered. **b**, Average amount of food consumed per day in vehicle and A20-dosed mice fed the GAN diet at the end of the study. **c**, Epididymal and inguinal white adipose tissue (eWAT, iWAT), brown adipose tissue (BAT), and heart body weight ratio in vehicle and A20-dosed mice fed the GAN diet (n = 4/5, *p<0.05, by Student’s t-test). **d-e**, Intraperitoneal (**d**) glucose tolerance test (GTT) and (**e**) Insulin tolerance test (ITT) was performed in these mice at week 36 (GTT) and week 38 (ITT) into the study. Data represent Mean ± SEM, ns= not significant.

